# Jaxley: Differentiable simulation enables large-scale training of detailed biophysical models of neural dynamics

**DOI:** 10.1101/2024.08.21.608979

**Authors:** Michael Deistler, Kyra L. Kadhim, Matthijs Pals, Jonas Beck, Ziwei Huang, Manuel Gloeckler, Janne K. Lappalainen, Cornelius Schröder, Philipp Berens, Pedro J. Gonçalves, Jakob H. Macke

## Abstract

Biophysiscal neuron models provide insights into cellular mechanisms underlying neural computations. However, a central challenge has been the question of how to identify the parameters of detailed biophysical models such that they match physiological measurements at scale or such that they perform computational tasks. Here, we describe a framework for simulation of detailed biophysical models in neuroscience—Jaxley—which addresses this challenge. By making use of automatic differentiation and GPU acceleration, Jaxley opens up the possibility to efficiently optimize large-scale biophysical models with gradient descent. We show that Jaxley can learn parameters of biophysical neuron models with several hundreds of parameters to match voltage or two photon calcium recordings, sometimes orders of magnitude more efficiently than previous methods. We then demonstrate that Jaxley makes it possible to train biophysical neuron models to perform computational tasks. We train a recurrent neural network to perform working memory tasks, and a feedforward network of morphologically detailed neurons with 100,000 parameters to solve a computer vision task. Our analyses show that Jaxley dramatically improves the ability to build large-scale data- or task-constrained biophysical models, creating unprecedented opportunities for investigating the mechanisms underlying neural computations across multiple scales.

## Introduction

Computational models are used to devise hypotheses about neural systems and to design experiments to investigate them. When building such models, a central question is how much detail they should include: Models of neural systems range from simple rate-based point neuron models to morphologically detailed biophysical neuron models [1–4]. The latter provide fine-grained mechanistic explanations of cellular processes underlying neural activity, typically described as systems of ordinary differential equations [5–10].

However, it has been highly challenging for neuroscientists to create biophysical models that can explain physiological measurements [10–13] or that can perform computational tasks [14–16]. It is hardly ever possible to directly measure all relevant microscopic properties of the system with sufficient precision to constrain all parameters directly, necessitating the use of *inference* or fitting-approaches to optimize free model parameters [17]. However, finding the right parameters for even a single neuron model with only a few parameters can be difficult [13, 18], and large-scale morphologically detailed biophysical network models may have thousands of free parameters. Therefore, neuroscien-tists work with simplified models that sacrifice biophysical detail for interpretability and computational efficiency (e.g., rate-based or leaky-integrate and fire neuron models, or models with simplified morphologies) [17, 19–21], or build biophysical simulations in a purely bottom-up way [11, 22, 23], without constraining the resulting network models by data or computational tasks as a whole.

Recently, in many domains of science such as particle physics, geoscience, and quantum chemistry, *differentiable*, GPU-accelerated simulators have enabled parameter inference for even complicated models using modern automatic differentiation techniques [24–28]. Such differentiable simulators make it possible to train simulators with gradient-descent methods from deep learning [29]: Backpropagation of error (‘backprop’) makes the computational cost of computing the gradient of the model with respect to the parameters independent of the number of parameters, making it possible to efficiently fit large models. In addition, GPU acceleration allows computing the gradient for many inputs (or model configurations) in parallel, which allows fitting simulations to large datasets with stochastic gradient descent [30]. Numerical solvers for biophysical models in neuroscience are used extensively, and several software packages exist [31, 32], in particular the commonly used N_euron_ simulation environment [33–35]. Yet, none of these simulators allows performing backprop, and currently used simulation engines are primarily CPU-based, with GPU-functionality only added post-hoc [36–38]. As a consequence, state-of-the-art methods for parameter estimation in biophysical neuron models are based on gradient-free approaches such as genetic algorithms [18, 39] or simulation-based inference [40], which do not scale to models with many parameters.

In principle, biophysical models of neurons are mathematically differentiable with respect to the parameters [41]. Thus, an efficient and scalable differentiable simulator would open up the possibility of optimizing such models with gradient descent. Such simulators would also need to handle the fact that the computation graph—a crucial ingredient for backpropagation of error—may be too large to keep in memory [42].

To close this gap, we developed a new framework for simulation and inference of (morphologically detailed) biophysical models in neuroscience, called Jaxley. Jaxley has in-built automatic differentiation capabilities and makes training efficient with native GPU-acceleration and just-in-time (JIT) compilation, building on computationally efficient engineering solutions developed in the machine learning community. This framework allows researchers to simulate large-scale biophysical models with thousands of parameters efficiently and even fit such models either to physiological data or to perform a computational task.

We demonstrate the power of this approach on a series of tasks covering different scales, data modalities, and levels of biophysical detail. Regarding fitting physiological data, we first show that we can efficiently recover the parameters of multicompartment models of single neurons based on experimental intracellular recordings or simulated voltage imaging recordings using gradient descent. In some cases, gradient descent is orders of magnitude more accurate and efficient than previous gradient-free methods. Second, we show that we can fit synaptic and cellular parameters of a hybrid model of the presynaptic circuit of a retinal ganglion cell to match dendritic calcium recordings in response to light stimuli. Regarding solving computational tasks, we show that we can train recurrent networks of Hodgkin–Huxley-type models (with detailed channel dynamics and multiple compartments) to solve working memory tasks. Finally, we build a large feedforward network with more than 800 cells, of which 64 are modelled at full morphological detail, and show that such a network can solve the classical MNIST task from machine learning without any additional nonlinearities. Our numerical experiments show that Jaxley is a flexible and easy-to-use simulation framework, running natively on CPUs, GPUs or TPUs and which, unlike previous neuroscience simulators, makes efficient automatic differentiation possible, unlocking new possibilities for data-driven biophysical simulations in neuroscience.

## Results

### Jaxley: A new toolbox for simulation and inference in neuroscience

Our goal was to optimize biophysical neuron models so that they either quantitatively fit data (e.g., match experimental recordings such as voltage or calcium measurements [11, 12, 18, 44]) or can achieve high performance in computational tasks such as short-term memory retrieval [45–47] or image classification [16, 48] (Fig. 1a). As the datasets that define these tasks are becoming increasingly complex, one has to adjust many (potentially thousands of) free parameters governing the behavior of ion channels (e.g., maximal conductance), synapses (e.g., synaptic conductance or time constant), or neural morphologies (e.g., radius or branch length). Inspired by the capabilities of deep learning to adjust millions (or even billions) of parameters given large datasets, we here suggest to adjust these parameters with gradient descent and to speed up training with GPUs.

**Figure 1.**
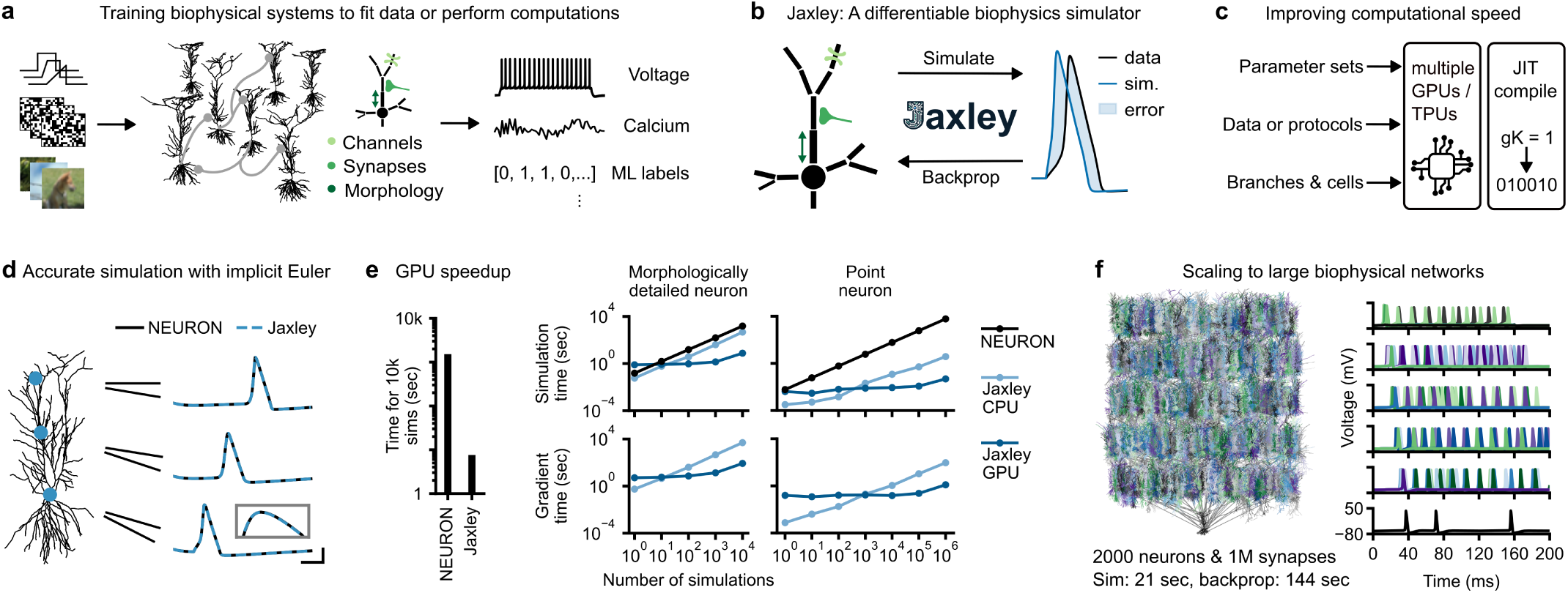
Differentiable simulation enables training biophysical neuron models. (a) Schematic of goal. Given input stimuli (left, step currents, or sensory stimuli encoded into input currents), we aim to train biophysically detailed neural networks (middle) either to match physiological recordings (right, voltage or calcium recordings), or to solve supervised machine learning tasks by predicting labels. To perform such tasks, we have to infer biophysical parameters such as channel conductances, synaptic conductances, or morphological parameters such as branch radii or lengths. (b) Schematic of method. Our simulator, Jaxley, allows to simulate biophysically detailed neural systems, and it can also compute the gradient of any cost function with respect to biophysical parameters via backpropagation of error. (c) Computational efficiency of Jaxley. Jaxley can parallelize simulations on (multiple) GPUs/TPUs across parameter sets, data (e.g., protocols), or branches and cells in a network. It can also just-in-time (JIT) compile code to further speed up simulation and training. (d) Simulated voltage traces at three locations, based on a reconstruction of a CA1 neuron [43] in response to a step current obtained with the Neuron simulator and with Jaxley. Inset is a zoom-in to the peak of the action potential. Scalebars: 3 ms and 30 mV. (e) Left: Time to run 10k simulations with Neuron on CPU and with Jaxley on GPU. Right: Simulation time (top) for the CA1 neuron shown in panel d and for a point neuron, as a function of number of simulations. Bottom: Same as top, for computing the gradient with backprop. (f) Jaxley scales to large networks. Jaxley can simulate and differentiate a biophysically detailed network built from reconstructions of CA1 neurons (left, network consists of 2,000 neurons and 1M synapses) in response to step currents to the first layer (voltage responses to the right). Runtimes were evaluated on an A100 GPU.

No current toolbox for biophysical simulation, however, allows to perform backprop as required for efficient gradient descent. Therefore, we built Jaxley, a new Python toolbox for simulation of biophysical neuron models. Jaxley allows to compute the gradient with respect to biophysical parameters with backprop, provides utilities to robustly perform gradient descent, and speeds up simulation and training using GPU acceleration and just-in-time (JIT) compilation (Fig. 1b). To achieve this, Jaxley implements numerical routines required for efficiently simulating biophysically-detailed neural systems, so-called implicit Euler solvers, in the deep learning Python framework JAX [49]. In particular, for multicompartment models, bespoke implicit solvers are key to accurate and stable simulation [50], but are not provided by any other toolbox for simulation of differential equations [51, 52]. Building on JAX, Jaxley inherits the capability to perform automatic differentiation, such that errors can be propagated back through the implicit Euler solvers, which makes it possible obtain the gradient with respect to any parameter.

Training biophysical models with gradient descent leads to instabilities resulting from parameters having different scales, networks having a large computation graph [42], loss surfaces being non-convex [53], and so on. Jaxley imple-ments methods that have been developed to overcome these specific issues in deep neural networks (Supp. Fig. S1). Jaxley also provides native support for parallelization: It implements the differential equations such that, for example, different branches of a single neural morphology can be solved in parallel on GPUs [36, 38, 54]. This results in large speed ups for cell and network simulation and differentiation. In addition, Jaxley can parallelize across stimuli or parameter sets, providing large speed-ups for datasets (via stimulus parallelization) or for parameter sweeps (via parameter parallelization, Fig. 1c). Finally, Jaxley just-in-time (JIT) compiles code, making it (at least) as fast as previous simulators which are written in compiled programming languages.

Taken together, the abilities of Jaxley to compute the gradient with backprop, to perform robust optimization with gradient descent, and to parallelize simulations across stimuli and cells on GPUs open up the possibility to train large biophysical networks with thousands of parameters. Furthermore, we designed Jaxley with a user-friendly interface, allowing neuroscientists to build biophysical models (e.g., for inserting recordings, stimuli, and channels into various branches or cells) and to use automatic differentiation and GPU parallelization. In a dedicated library open to the community, it also implements different types of connectivity structures (e.g., dense or sparse connectivity), a growing set of ion channel models, and utilities that are required to perform robust training with gradient descent (e.g., parameter transformations, multi-level checkpointing [55], specific optimizers for non-convex loss surfaces [56]). Jaxley is fully written in Python, which will make it easy for the community to use and to add functionality to it^1^.

### Jaxley is accurate, fast, and scalable

We benchmarked the accuracy, speed, and scalability of Jaxley for simulation of biophysical models. First, we evaluated the accuracy of Jaxley and created biophysically-detailed multicompartment models of a CA1 pyramidal cell from rat hippocampus [43, 57] and of four layer 5 neurons from the mouse visual area from the Allen Cell Types Database [58]. Every model contained sodium, potassium, and leak channels in all branches. We stimulated the soma and recorded the voltage at three locations across the dendritic tree. Jaxley matched the voltages of the Neuron simulator at sub-microsecond and sub-millivolt resolution (Fig. 1d). Across all five cells, input currents with ten different amplitudes, and three recording sites, the deviation of spike time between the Neuron simulator and Jaxley was at most 0.05 ms and the difference in spike amplitude was at most 0.02 mV (Supp. Fig. S2).

Next, we evaluated the simulation speed of Jaxley on CPU and GPU. We simulated the above described CA1 cell, as well as a single compartment model, for 20 ms. To demonstrate the parallelization capabilities of Jaxley, we also evaluated the runtimes for running multiple simulations in parallel with different parameter sets. On GPU, Jaxley was much faster for large systems or many simulations, with a speed up of around two orders of magnitude (Fig. 1e). For single compartment neurons, Jaxley could parallelize the simulation of up to 1 million neurons, thereby allowing fast parameter sweeps. On CPU, Jaxley was similar to Neuron in speed for multicompartment models and was faster than Neuron for single compartment models because Jaxley can vectorize code, which avoids slow for-loops across parameter sets.

We then evaluated the computational cost of computing the gradient with Jaxley (Fig. 1e, bottom). For back-propagation, the forward pass must be stored in-memory, which can easily correspond to terabytes of data for large neural systems. To overcome this, Jaxley implements multi-level checkpointing [55], which reduces memory usage by strategically saving and recomputing intermediate states of the system of differential equations. Overall, we found that computing the gradient was slightly more expensive than the simulation itself: Depending on the simulation device (CPU/GPU/TPU), the number of simulations, the number of branches and channels, the simulated time, and the loss function, computing the gradient was between 3 and 20 times more expensive than running the simulation itself (Fig. 1f, bottom) [59].

Finally, we show that in addition to parallelizing across parameters (or across stimuli), Jaxley can parallelize across branches or compartments in a network, allowing to simulate and differentiate large networks of biophysical neuron models. To demonstrate this, we built a multi-layer neural network consisting of 2,000 morphologically detailed neurons with Hodgkin–Huxley dynamics, connected by 1 million biophysical synapses (3.92 million differential equation states in total, Fig. 1f). On a single A100 GPU, Jaxley computed 200 ms (i.e., 8,000 steps at Δ*t* = 0.025 ms) of simulated time in 21 seconds. We then used backprop to compute the gradient with respect to all membrane and synaptic conductances in this network (3.2 million parameters in total), which took 144 seconds. Estimating the gradient with finite differences—as would be required for packages that do not support backprop—would take more than two years (3.2 million forward passes, 21 seconds each).

### Fitting biophysical single neuron models to intracellular recordings

Having demonstrated the accuracy and speed of Jaxley, we applied it to a series of tasks which demonstrate how Jaxley opens up new opportunities for building task-or data-driven biophysically-detailed neuroscience models in a range of scenarios. We show that Jaxley can fit biophysical models to physiological measurements such as voltage and calcium recordings, sometimes vastly more accurately and efficiently than gradient-free methods, and we demonstrate that Jaxley allows to fit large-scale biophysical networks with up to 100k parameters to computational tasks such as memory retrieval or image recognition.

As a first proof-of-principle, we applied Jaxley to fit single neuron models with few parameters. As we will show, even in these models in which gradient-free methods such as genetic algorithms excel and are used extensively, gradient descent can be competitive and sometimes even outperform state-of-the-art genetic algorithms. We built a biophysical neuron model based on a reconstruction of a layer 5 pyramidal cell (L5PC) (Fig. 2a). The model had nine different channels in the apical and basal dendrite, the soma, and the axon [39], with a total of 19 free parameters, including maximal channel conductances and dynamics of the calcium pumps. We learned these parameters from a synthetic somatic voltage recording given a somatic step current stimulus with a known set of ground-truth parameters (Fig. 2b, top).

**Figure 2.**
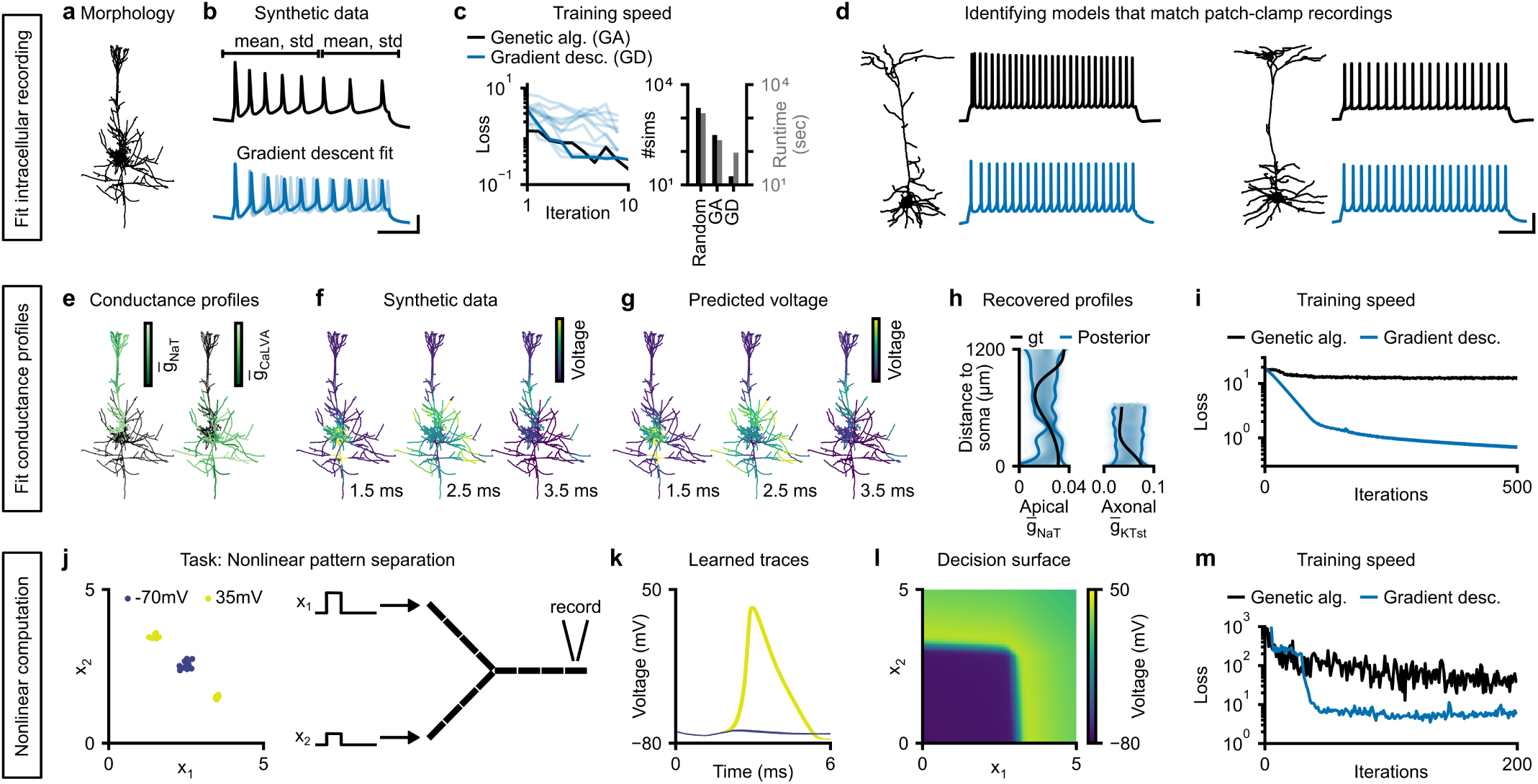
Inferring single-neuron models with gradient descent. **Task 1** (a) We optimize 19 parameters of a biophysical neuron model which is based on the morphology of a layer 5 pyramidal cell (L5PC) morphology. (b) Top: Synthetic somatic voltage recording (black) and windows that are used to compute summary statistics (top). Bottom: Fits obtained with gradient descent. Best fit in dark blue, fits from five best independent runs in light blue. Scalebars: 20 ms and 30 mV. (c) Left: Loss value of individual gradient descent runs (light blue), their minimum (dark blue), in comparison to the minimum loss across ten genetic algorithm runs (black). Right: Average number of simulations and runtime required to find a set of parameters with loss smaller than 0.55 with random sampling, genetic algorithm, and gradient descent. (d) We also fit morphologically detailed models to patch-clamp recordings (black) in response to step currents from the Allen Cell Types Database. Gradient descent fit in blue. Additional models in Supp. Fig. S3. Scalebars: 200 ms and 30 mV. **Task 2** (e) We optimize conductance profiles of the same L5PC morphology, leading to 1390 parameters. Synthetic ground-truth conductance profiles vary as a function of distance to the soma. (f) Simulated voltages given the synthetic conductance profile after 1.5, 2.5, and 3.5 ms. (g) Predicted voltages of gradient descent fit closely match synthetic observation. (h) Ground truth conductance profile as a function of distance from the soma (black) and 90 % confidence interval obtained with multi-chain gradient-based Hamiltonian Monte-Carlo. (i) Loss as a function of the number of iterations for gradient descent and genetic algorithm. **Task 3** (j) We optimize parameters of a simplified morphology with twelve compartments (right) to solve a nonlinear pattern separation task (left). (k) Voltage traces of model found with gradient descent. (l) Decision surface of the model reveals nonlinear single-neuron computation. (m) Minimum loss across ten independent runs as a function of the number of iterations for gradient descent and genetic algorithm.

We used gradient descent to identify parameter sets which minimize the mean absolute error to summary statistics of the voltage trace. Since gradient descent requires differentiable summary statistics, but commonly used summary statistics of intracellular recordings—such as spike count—can be discrete or non-differentiable, we used mean and standard deviation of the voltage in two time windows [40]. Starting from randomly initialized parameters, gradient descent required only nine steps (median across ten runs) to find models whose voltage traces are visually similar to the observation (Fig. 2b, bottom). A state-of-the-art indicator-based genetic algorithm (IBEA) required similarly many iterations, although each iteration of the genetic algorithm used ten simulations. As a consequence, gradient descent required almost ten times fewer simulations than the genetic algorithm, and, despite the additional cost of backpropagation, found good parameter sets in less runtime than the genetic algorithm on CPU (Fig. 2c).

We then employed gradient descent to identify parameters that match patch-clamp recordings from four cells from the Allen Cell Types Database which had an axon initial segment reconstructed (because many of our parameters were axonal, Fig. 2d, top) [58]. We again defined windows of the voltage for summary statistics, inserted the same set of ion channels, and used the same set of free parameters (see Methods). Due to the length of the recordings (1 s vs 100 ms in the synthetic experiments), this was a much more challenging problem as it could lead to exploding or vanishing gradients. Despite this, gradient descent found parameter sets whose voltage traces closely resembled experimental recordings (Fig. 2d, bottom, additional fits in Supp. Fig. S3). For both models, gradient descent was more simulation efficient than the genetic algorithms and had similar runtime (Supp. Fig. S4). Overall, these results demonstrate the ability of gradient descent to fit biophysical models to intracellular recordings, being competitive with state-of-the-art genetic algorithms even on tasks for which those have been extensively optimized.

### Fitting biophysical single neuron models with many parameters

How does gradient descent scale to models with large numbers of parameters? We demonstrate here that, in contrast to genetic algorithms, gradient descent allows to optimize a single neuron model with 1390 parameters.

We used the above described model of a L5PC with a diverse set of active conductances. Unlike in the above experiments, we fit the maximal conductance of ion channels in every branch in the morphology [60], thereby allowing to model effects of non-uniform conductance profiles [61, 62]. This increased the number of free parameters to 1390. To generate a synthetic recording, we assigned a different maximal conductance to each branch (sampled from a Gaussian process, see Methods), depending on the distance from the soma (Fig. 2e). We recorded the voltage at every branch of the model in response to a 5 ms step current input (Fig. 2f). Experimentally, such data could be obtained, for example, through voltage imaging [63].

We employed gradient descent to identify parameters that match this recording, with a regularizer that penalizes the difference between parameter values in neighboring branches, thereby targeting conductance profiles which vary smoothly across the dendritic tree [62, 64]. Despite the large number of parameters, gradient descent found a parameter set whose voltage response closely matched the observed voltage throughout the dendritic tree (Fig. 2g). To understand how much the whole cell voltage recording constrains the parameters [41], we used Bayesian inference (implemented with gradient-based Hamiltonian Monte–Carlo, details in Methods) to infer an ensemble of parameter sets all of which match the observed voltage. The resulting ensemble revealed regions along the dendritic tree at which the conductance profile was strongly constrained by the data (e.g., the transient sodium channel, Fig. 2h, left, e.g., around 400 µm). The parameter ensemble also revealed conductance profiles (e.g., axonal low-voltage activated calcium channel) which were only weakly constrained by the data, suggesting that diverse values of these parameters could lead to the desired activity (Fig. 2h, right, all posterior marginals in Supp. Fig. S5) [65]. Finally, we compared our method with an indicator-based genetic algorithm [18] and, as expected, we found that access to gradients leads to better convergence: While gradient descent converged to values of low loss within 100 iterations, the genetic algorithm had two orders of magnitude higher loss even after 500 iterations (Fig. 2i).

### Nonlinear single neuron computation

In the two previous tasks, we showed that Jaxley can learn model parameters such that simulations match recordings, and that it can do so as efficiently as state-of-the-art methods for small neural models and much more efficiently for large ones. Next, we demonstrate that gradient descent—commonly used to fit deep neural networks—can also be used to train single neurons to perform computational tasks.

We trained a single neuron model to solve a nonlinear pattern separation task. While point neurons cannot solve such tasks (because they linearly sum their inputs), it has been suggested that morphologically-detailed neurons might solve nonlinear pattern separation tasks in their dendritic tree [66–69]. While it has been demonstrated extensively that single neuron models respond nonlinearly to inputs [70–72], it has so far been difficult to train biophysically-detailed neurons on a particular task. Here, we show that stochastic gradient descent allows to train single neuron models with dendritic nonlinearities to perform nonlinear computations.

We defined a simple morphology consisting of a soma and two dendrites, and inserted sodium, potassium, and leak channels into all neurites of the cell. We then learned ion channel densities as well as length, radius, and axial resistivity of every compartment (72 parameters in total) for the neuron to have two outputs depending on the input: low somatic voltage (−70 mV), when both dendrites were stimulated with step currents of intermediate strength; high somatic voltage (35 mV), when one of the dendrites was stimulated strongly and the other one weakly (Fig. 2j). Therefore, the two classes were not linearly separable, requiring the neuron to perform a nonlinear computation.

After training the parameters with gradient descent, we found that the cell indeed learned to perform this task and spiked only when one dendrite was stimulated strongly (Fig. 2k), effectively having a nonlinear decision surface (Fig. 2l). We again compared our method to an indicator-based genetic algorithm and found that our method finds regions of lower loss more quickly than genetic algorithms (Fig. 2m).

Overall, these results show that gradient descent performs better than gradient-free methods in models with many parameters, opening up possibilities for studying at scale biophysical mechanisms throughout the full neuronal morphology.

### Hybrid retina model of dendritic calcium measurements

So far, we have learned parameters of single neuron models using small datasets consisting of few stimulus/response pairs. Many models of neural systems, however, consist of multiple neurons potentially modelled at varying levels of detail, and datasets can contain thousands of stimulus/response pairs [16, 44, 74, 75]. Using a network model of the mouse retina, we demonstrate that Jaxley allows to simultaneously infer cell-level and network-level parameters, and that it can identify parameters such that model simulations match datasets consisting of thousands of stimulus/response pairs.

As an example, we consider transient Off alpha retinal ganglion cells in the mouse retina, which show compart-mentalized calcium signals in their dendrites in response to visual stimulation [73]. To understand the mechanistic underpinning of this behaviour, we built a hybrid model with statistical and mechanistic components: We modelled photoreceptors as convolution with a Gaussian filter, bipolar cells as point neurons with a nonlinearity [74], and a retinal ganglion cell (RGC) as a morphologically detailed biophysical neuron, with six different ion channels [76] distributed across its soma and dendritic tree (Fig. 3a). In order to model the recording of calcium signals, we convolved the intracellular calcium (from the calcium channel of the model) with a calcium kernel (Fig. 3b).

**Figure 3.**
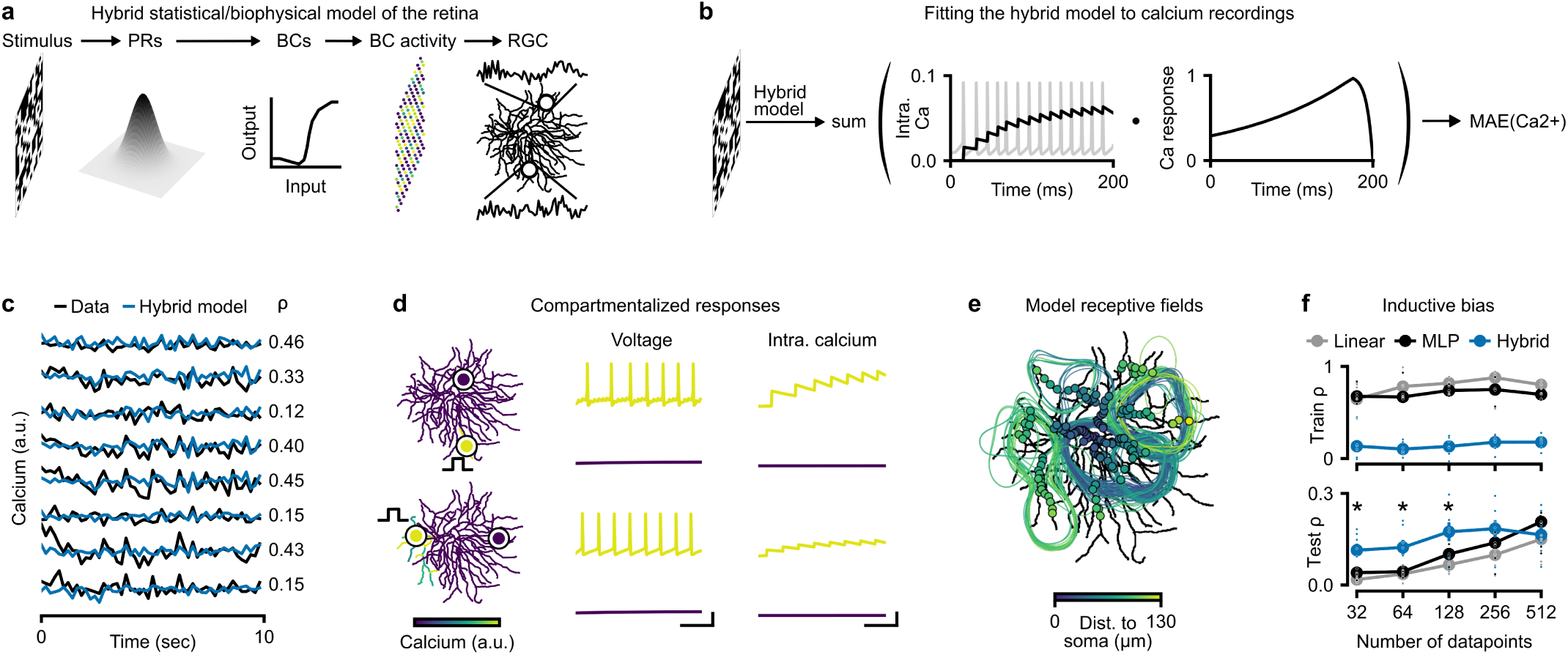
Hybrid model of the calcium responses of a retinal ganglion cell. (a) Schematic of experimental setup and hybrid model. A binary noise image (15 *×* 20 pixels) was presented to the retina of mice and dendritic calcium recordings were obtained with two-photon imaging [73]. We modelled photoreceptors (PR) as linear Gaussian filters, bipolar cells (BC) as point neurons with a non-linearity, and a retinal ganglion cell at full morphological detail with a variety of ion channels. (b) Schematic of training procedure and loss function. We stimulated the hybrid model with the noise image for 200 ms and recorded intracellular calcium concentration across all recording sites. We convolved this concentration with a calcium kernel and used the output at the last time step (200 ms) as input to a mean absolute error cost function to the experimental recording. (c) Measured and model-predicted calcium response across 50 noise images (200 ms each). (d) Left: Calcium response (colormap) of the trained hybrid model to a step-current to a single branch indicated by step current sketch. Middle: Voltage activity of the model at two branches, one at the stimulus site and one at a distant branch. Right: Intracellular calcium concentration in the same two recording sites. Scalebars: 50 ms, 30 mV, 0.025 mM. (e) Receptive fields of the hybrid model obtained in response to 1024 noise stimuli. (f) Pearson correlation coefficient between experimental data and model for train (top) and test (bottom) data, for a linear network, a multi-layer perceptron, and the hybrid model. Error bars show standard-error of mean over seven datasets (see Methods). Asterisk denotes statistically significant difference between mean correlations of hybrid model and MLP (one-sided t-test at p<0.05).

Using Jaxley, we trained the hybrid model to predict dendritic calcium given checkerboard noise stimuli. The dataset consisted of 15,000 image-calcium pairs, with each image being presented for 200ms. We learned synaptic conductances from the bipolar cells onto the RGC (287 synaptic parameters), the radii of the branches of the retinal ganglion cell, the axial resistivities, as well as the somatic and basal membrane conductances (320 cell parameters). After training, we evaluated the trained model on a held-out test dataset. The model had a positive Pearson correlation coefficient with the experimental recording on 146 out of 147 recordings sites, with an average correlation of 0.25, and a maximum of 0.51 (Fig. 3c).

Next, we tested whether the hybrid model was able to predict experimentally measured phenomena which the model had not been directly trained on. To test whether our model showed the same compartmentalized calcium responses as the measurements [73], we stimulated the trained model at a distal branch and recorded the model calcium response across all branches of the cell (Fig. 3d, left). We found that the calcium signal in response to local stimulation did not propagate through the entire cell, demonstrating a compartmentalized response of the model. While the neurons often exhibited high firing rates near the stimulation site in the dendrite (Fig. 3d, middle, [77, 78]), branches that were far away from the stimulation site showed no response (Fig. 3d, right). To further demonstrate the compartmentalized response of the hybrid model, we computed the receptive fields of all experimental recordings sites in response to noise images (Fig. 3e). The model receptive fields did not cover the entire cells and were roughly centered around the recording locations, qualitatively matching the receptive fields obtained from experimental measurements [73].

The mechanistic components of the model, including the anatomical structure, provide an inductive bias. Therefore, we investigated whether this inductive bias of the hybrid model could lead to better generalization to new data, especially when training data is scarce. We trained the hybrid model on reduced datasets of recordings from a single calcium scanfield and on a reduced set of images, and compared its performance to a linear model and a two-layer perceptron trained on the same datasets (Fig. 3f). For all models, we performed early stopping based on a validation set and evaluated the final performance on a held-out test set of 512 images. While the linear model and the perceptron performed better than the hybrid model on training data, the hybrid model performed better on held-out test data, when little training data was available. These results indicate that the inductive bias brought by the hybrid model effectively can limit the amount of overfitting in the model, and allow the model to have higher generalization performance than unconstrained artificial neural networks, at least in small data-regimes. These results suggest that hybrid components could be used as regularizers for deep neural networks models of neural systems [79–83].

Our results demonstrate that gradient descent allows to fit networks of biophysical neurons to large calcium datasets and allows to simultaneously learn cell-level and network-level parameters. The resulting models reproduce several experimental measurements and exhibit an inductive bias that leads to improved generalization performance on small datasets.

### Biophysical recurrent network models can be trained to solve working memory tasks

To understand how computations are implemented in neural circuits, computational neuroscientists aim to train models to perform tasks [16, 45–48]. In particular, recurrent neural networks (RNNs) have been used to form hypotheses about population dynamics underlying cognition [15, 19, 45–47, 84, 85]. Typically, such RNNs consist of point neurons with rate-based or simplified spiking dynamics [42, 86], which prevents studying the contribution of channel dynamics or cellular processes [87]. We here show how Jaxley makes is it possible to train biophysical models of neuronal networks to perform such tasks.

We implemented in Jaxley an RNN consisting of Hodgkin–Huxley-type neurons with a simplified apical and basal dendrite, with each neuron equipped with a variety of voltage-gated ion channels [11]. We sparsely connected the recurrent network with conductance-based synapses [88] and obtained the outputs from passive readout units (Fig. 4a).

**Figure 4.**
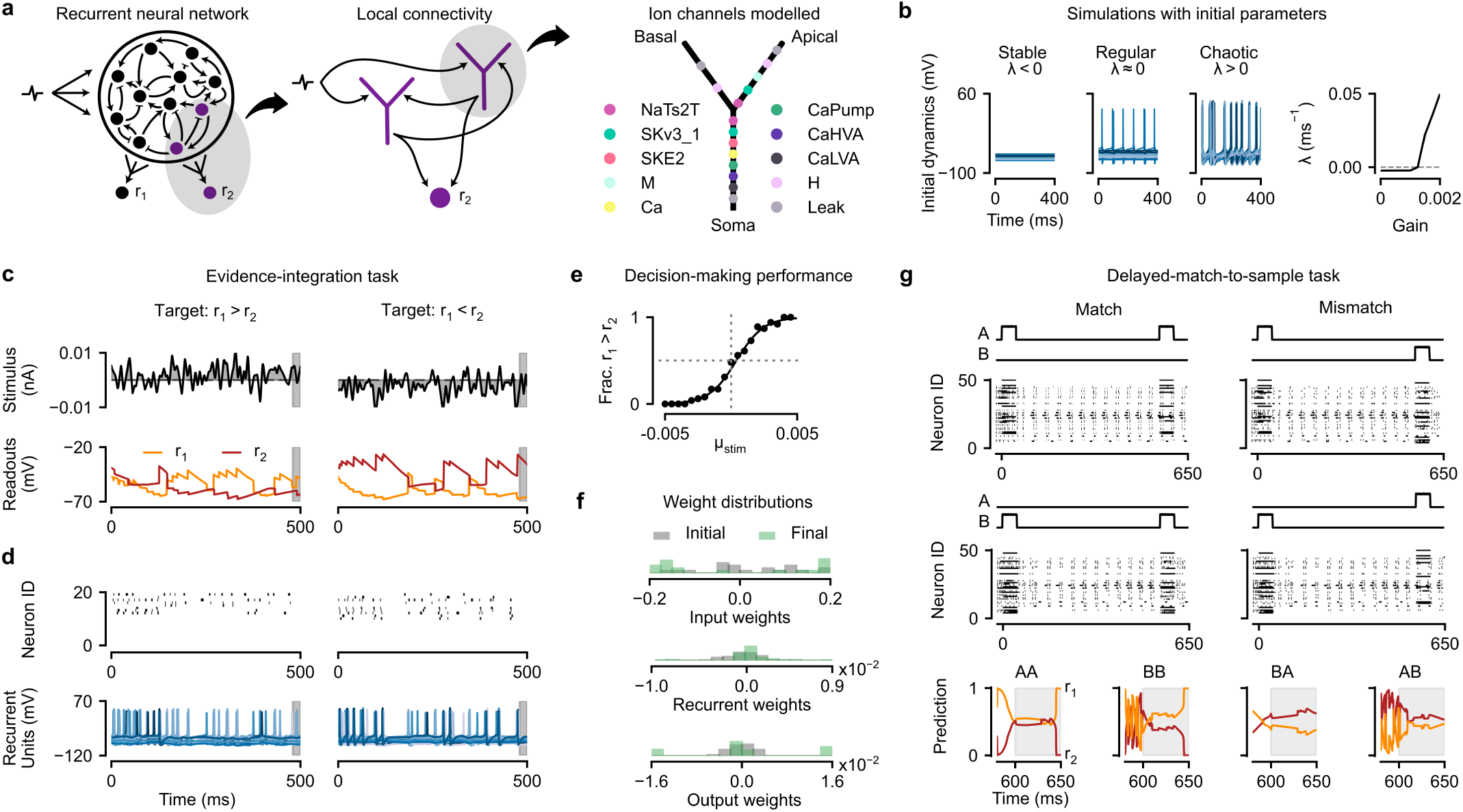
Recurrent neural network models with Hodgkin–Huxley-type neurons perform working memory tasks. (a) Left: Schematic of the RNN, with two recurrently connected neurons and one readout neuron highlighted. Neurons get stimulated at their basal dendrites and have recurrent synapses from somata to apical dendrites. Right: The ion channels modelled in each neuron. (b) Dynamics (left) and associated Lyapunov exponents (right) of the recurrently-connected neurons before learning the parameters and without any stimulus. The dynamics vary as a function of the scale (or ‘gain’) of the recurrent synaptic conductances. **Task 1** (c) Evidence integration task. We stimulated an RNN with a random Gaussian noise stimulus (top). After training, the RNN successfully learns to indicate whether the summed inputs are greater than zero (left) or less than zero (right), as indicated by voltage traces of the two readout neurons during the response period (grey). (d) Raster plots (upper) and voltage traces (lower) of the recurrently-connected neurons encoding the stimulus. (e) Psychometric curve showing the fraction of times the RNN reported the summed input to be greater than zero (*r*_1_ *> r*_2_), as a function of the stimulus mean (which modulates task difficulty). (f) Histogram of initial and trained input, recurrent, and output weights of the network. **Task 2** (g) Delayed-match-to-sample task. The RNN is presented with two stimuli (A/B) separated by a delay, and was trained to indicate whether the two stimuli ‘match’: The match case is where A or B contain both input step currents, and the mismatch case is where A and B each contain one input step current. Raster plots of the network activity, when presented with each of the four scenarios. Bottom row: Readout neuron prediction in the match (left, *r*_1_ *> r*_2_) and mismatch (right, *r*_2_ *> r*_1_) cases.

We first investigated the dynamics in this biophysical RNN before training. As with rate-based RNNs, these dynamics were strongly dependent on a global scaling factor (called ‘gain’) of all recurrent synaptic maximal conductances [89]. Our RNN transitioned from a stable to a chaotic regime when the gain was increased, with an intermediate region, where networks displayed regular firing (Fig. 4b, left). The ability of Jaxley to perform automatic differentiation allowed us to quantify the stability of networks by numerically computing Lyapunov exponents [90, 91] (Supp. Fig. S7).

When we increased the gain of the synapses, the maximal Lyapunov exponent transitions from a value smaller than 0, corresponding to stable dynamics, to a value larger than 0, corresponding to a chaotic system, where nearby trajectories diverge (Fig. 4b, right). This indicates that, upon parameter initialization, biophysical RNNs have similar dynamical regimes as rate-based RNNs.

We then trained the biophysical RNN to perform two working memory tasks, starting with a perceptual decision-making task requiring evidence integration over time [45, 47, 92–94]. We built a network of 20 recurrent neurons and stimulated each recurrent neuron with a noisy time series with either positive or negative mean value (Fig. 4c, top, see Methods). We trained input weights, recurrent weights, and readout weights (109 parameters) such that the network learned to differentiate between positive and negative stimuli during a response period after 500 ms. Despite the long time horizon of this task (500 ms, corresponding to 20k time steps of the simulation), gradient descent found parameters such that the RNN was able to perform the task (Fig. 4c, bottom), with the voltage in the readout neurons differentiating the input means with 99.9 % accuracy across 1,000 trials. The trained network showed sparse spiking activity with typically less than one third of neurons being active for a particular stimulus (Fig. 4d, top). Furthermore, the dynamics of the trained network without stimulus input were chaotic (Lyaponuv exponents close to 0, *λ* = 2 · 10^−3^, Fig. 4d, bottom), even though the initial dynamics were not (*λ* = −1 · 10^−5^), consistent with previous studies linking the regime close to ‘the edge of chaos’ to optimal computational performance [95, 96].

Next, we evaluated the generalization abilities of the trained biophysical RNN. We varied the mean value of the positive and negative stimuli and found that the relationship between average response and stimulus mean closely resembled the well-known sigmoidal psychometric curve of decision making, where the network more often failed when the stimulus had a lower signal-to-noise ratio (Fig. 4e). We also tested whether the network generalized in time and found that, despite having only been trained on tasks of 500 ms duration, the RNN could successfully solve evidence integration tasks of up to 3 seconds (Supp. Fig. S6). To solve these evidence integration tasks, some input, recurrent and output weights were pushed toward zero and the remaining weights were pushed toward their positive and negative constraints during training (Fig. 5f).

**Figure 5.**
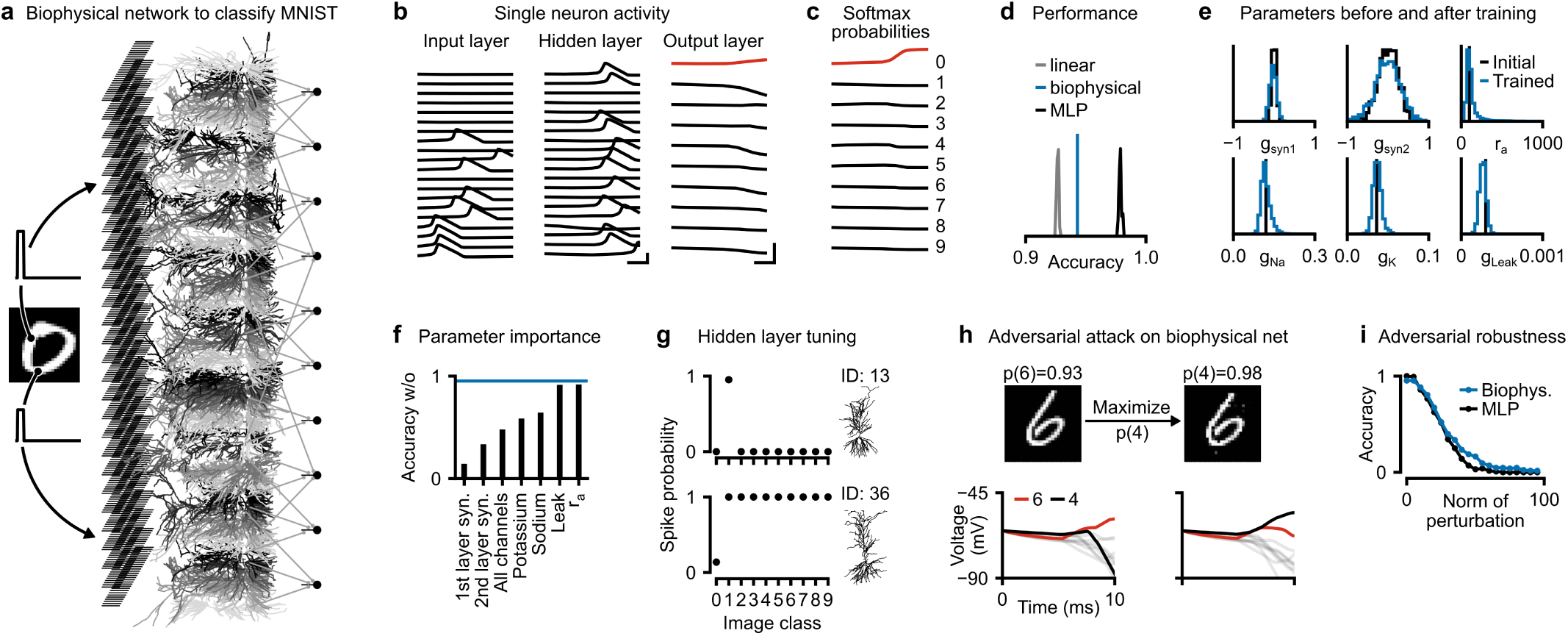
Training biophysically detailed networks to solve computer vision tasks. (a) We trained a biophysical network consisting of 28 *×* 28 input neurons, 64 morphologically detailed hidden neurons, and ten output neurons. We stimulated input neurons with step currents corresponding to MNIST pixel values and used voltages of the somata of output neurons after 10 ms as prediction of class membership. (b) Voltages measured at the somata of neurons in the trained network in response to an image labelled as ‘0’. Red color in the output layer indicates the image label. Scalebars: 2 ms and 80 mV. (c) Softmax probabilities computed from output voltages. (d) Histograms of test set accuracy of 50 linear networks (gray), 50 multi-layer perceptrons with 64 hidden neurons and ReLU activations (black), and the biophysical network (blue). (e) Histogram of parameters before (black) and after (blue) training. Training the network only leads to subtle shifts in parameter distributions. (f) Test set accuracy for trained network when subsets of parameters are reset to their initial value. Blue line is the full trained network. (g) Fraction of images that trigger a spike in two example hidden neurons across images from different classes. Top neuron has acquired ON-tuning for the digit ‘1’, bottom neuron has OFF-tuning for the digit ‘0’. (h) Adversarial attack on the biophysical network. The trained biophysical network correctly classifies the image as a six with high confidence. After modifying the image with an adversarial attack, the image remains similar but the biophysical network confidently predicts a four. Bottom shows corresponding voltage traces of the output neurons (output neurons corresponding to other digits in gray). (i) Accuracy across 128 test set examples, as a function of the norm of the adversarial image perturbation.

We next employed the RNN to solve a more challenging working memory task, a delayed match to sample task, where the RNN had to maintain information over an extended period of time [46, 47, 86, 97]. We trained the RNN to classify patterns, consisting of two step current inputs with a delay between them, into matching (same identity of the inputs) or non-matching (different identity of the inputs), i.e., a total of 4 input patterns (Fig. 4g, top). A central challenge in this task was that the network needed to memorise the identity of the first input within its dynamics for a delay period of 500 ± 50 ms until the second input was provided. We employed curriculum learning to solve this task: Starting from shorter delay periods of 100 ± 50 ms, we increased the delay period during training in steps of 100 ms [98]. By training input, recurrent, and readout weights, as well as synaptic time-constants of a network with 50 recurrent neurons (542 parameters), we found parameter sets which solved the task and correctly classified all four patterns (Fig. 4g, bottom).

Previous studies found that rate-based RNNs could learn working memory tasks either using transient coding or stable attractors [99–101]. We investigated which of those two mechanisms the biophysical RNN used to successfully maintain the stimulus identify during the delay period. To do so, we inspected the population dynamics of the trained biophysical RNN. The network had distinct responses to the identity of the first stimulus (Fig. 4g) and only when running the simulation much longer than the delay period the dynamics relaxed to the same attractor (Supp. Fig. S8). This suggests that the network used a form of transient coding to solve the task (depending on initialisation and training setup, other solutions might be possible [101]).

Overall, these results demonstrate that gradient descent allows training RNNs with biophysical detail to solve working memory tasks. This will allow a more quantitative investigation of the role of cellular mechanisms contributing to behavioral and cognitive computations.

### Training biophysical networks with 100k parameters on large datasets

Finally, we show that gradient descent allows to train large biophysical models with thousands of cellular level and network-level parameters on machine learning-scale datasets to solve classical computer vision tasks like image recognition.

As a demonstration, we implemented a feedforward biophysical network model in Jaxley and trained it to solve the classical MNIST task, without artificial nonlinearities such as ReLU activations. The network had three layers: The input and output layers consisted of neurons with ball-and-sticks morphologies and the hidden layer consisted of 64 morphologically detailed models obtained from reconstructions of CA1 cells (Fig. 5a) [43, 54]. The input and hidden layer had active ion channels (sodium, potassium, leak) in all branches, thereby permitting nonlinear computation as spike/no-spike decisions and allowing the network to have nonlinear responses. The output neurons only contained a leak channel and integrated hidden-layer activity. The network was interconnected by biophysical synapses [88] that could be excitatory or inhibitory. We trained sodium, potassium, and leak conductances of every branch in the circuit (55k parameters), as well as all synaptic weights (51k parameters).

We simulated the network for 10 ms, as this was the time it took for the stimulus to propagate through the network. After training with stochastic gradient descent (see Methods), upon being stimulated with a ‘0’ digit, the network propagated spikes through the first two layers, such that the somatic voltage of the output neuron corresponding to ‘0’ had high voltage after 10ms (Fig. 5b). When passed through a softmax, the output neuron activations indeed indicated a high probability for the digit ‘0’ (Fig. 5c). The network achieved an accuracy of 94.2 % on a held-out test dataset, which is higher than a linear classifier, demonstrating that the biophysical network successfully uses its nonlinearities to improve classification performance. The biophysical network, however, performed slightly worse than a multi-layer perceptron with ReLU nonlinearities, suggesting that the spike/no-spike nonlinearitites are either more difficult to train than ReLU nonlinearities, or that the (binary) spike/no-spike representations lead to lower bandwidth than graded ReLU activations (Fig. 5d).

How do the learned parameters of the biophysical network contribute to its ability to classify MNIST digits? Surprisingly, we found that the ranges of the trained synaptic parameters were roughly similar to the ranges of the untrained network and that the membrane channel conductances were roughly centered around their initial value (Fig. 5e). This does, however, not mean that the learned values of these parameters do not contribute to the learned network dynamics: When resetting subsets of parameters to their initial value, classification dropped substantially, sometimes reducing classification performance to chance level accuracy of 10 % (Fig. 5f). Resetting some parameters (e.g., leak conductances or axial resistivities) had a weaker but still substantial effect on performance (reduced accuracy to 91 %). Overall, although the average values of parameters remained roughly unchanged from initialization, the respective tuning of virtually all parameters in the system, including ion channel conductances, contributed to the learned network dynamics and classification performance. This indicates that biophysical simulations built purely from the aggregate statistics of measured parameter values could not be sufficient for the models to develop the ability to perform tasks [11].

We then investigated which features the morphologically detailed hidden neurons were tuned to. Across 2560 images, we evaluated the fraction of images from each digit to trigger a spike in each hidden neuron. We found several neurons that were strongly tuned to individual digits: For example, we found a neuron which spiked for almost any ‘1’ in the dataset, but rarely spiked for images of other digits in the dataset, suggesting that this cell had ON-tuning for the ‘1’ digit, just one set of synapses away from the input layer (Fig. 5g, top). We also found neurons which exhibited OFF-tuning for some digits: For example, a hidden neuron spiked for any image which did not show a ‘0’, but spiked only for a fraction of ‘0’ images (Fig. 5g, bottom, tuning of all hidden neurons in Supp. Fig. S9). Neither of these neurons were tuned to individual digits before training (Supp. Fig. S10). These findings show that the trained biophysical network had hidden neurons strongly tuned to interpretable features, which emerged from training the network to solve a task.

Finally, efficient access to gradients does not only allow training biophysical models to match physiological data or perform a task, but it also opens up possibilities to perform new kinds of analyses on them, like evaluating adversarial robustness [102]. To illustrate this, we stimulated the biophysical network with an image showing a ‘6’, and then altered minimally the image with gradient descent such that the network would classify it as a ‘4’ (Fig. 5h). While the biophysical network classified the initial image as a ‘6’ with high confidence, the perturbed image was classified as a ‘4’—despite only weak and barely noticeable changes to the image. We compared the adversarial robustness of the biophysical network to a trained multi-layer perceptron (MLP) with ReLU activations and the same number of hidden neurons. We found that the two networks are similarly vulnerable to adversarial attacks, and any improvement in adversarial robustness was comparably small, in contrast to previous studies suggesting that biophysical networks could largely improve adversarial robustness [54].

Overall, these results demonstrate that the ability of Jaxley to perform backprop in biophysical models can be used to train large-scale networks at full biophysical detail. In particular, backprop overcomes virtually any computational limit on the number of parameters of biophysical networks that can be included in the optimization, thereby opening up new possibilities for task-trained biophysical models.

## Discussion

Backpropagation of error (‘backprop’) and computational frameworks which provide efficient, scalable, and easy-to-use implementations, have been key to to the deep learning revolution. They have made it possible to efficiently optimise even very big systems with gradient descent. This, in turn, has made it possible to fully optimise entire machine learning workflows, and minimised the need for hand-designing (‘feature engineering’) its components. Several scientific disciplines are now adopting such ‘differentiable programming’ approaches in which entire pipelines are implemented as differentiable simulators and can therefore be optimised or fit to data [24–28, 103, 104]. However, for biophysical neuron models, currently used tools [31, 32, 105, 106] do not allow automatic differentiation. We here presented Jaxley, a new computational framework for differentiable simulation of neuroscience models with biophysical detail. Unlike previous biophysical simulation toolboxes, Jaxley can perform automatic differentiation through its differential equation solver, thereby enabling backprop to compute the gradient with respect to virtually any biophysical parameter. We demonstrated that gradient descent allows to fit biophysical models of neural dynamics to large datasets of experimental voltage and calcium recordings and that it enables training biophysical models to perform physiologically meaningful computations, with as many as 100k parameters.

We expect that Jaxley will enable a range of new investigations in neuroscience: First, it will make it possible to efficiently optimize detailed single-cell models. This will allow insights into cellular properties across cell types and their relationship with, for example, transcriptomic measurements [107–109], as well as into the contribution of dendritic processing to neural computations [70, 72, 110–113].

Second, it will facilitate creating large-scale biophysical network models. Such network models [11, 22, 114] have so far primarily been built in a bottom-up fashion: Models of single biophysical neurons are individually fit to electrophysiological recordings using genetic algorithms, with minimal adjustments of the network as a whole, and computational properties are thought to emerge through this process. This approach has been successful in creating models which can reproduce aggregate statistics of neurophysiological measurements such as local field potentials [11, 115–117]. However, it is unlikely that such networks will be able to explain behavioural or cognitive computations. A prerequsiste for this would be that these networks can perform relevant computational tasks such as image recognition or working memory tasks. However, a central impediment has been the lack of computational frameworks that allow to adjust (potentially thousands of) parameters of these models accurately and efficiently [10, 13, 18]. Inspired by the optimisation methods for deep neural networks which can fit millions of parameters with backprop and are accelerated by GPU parallelization, Jaxley opens up possibilities to train parameters of large-scale biophysical network models with gradient descent. We note that, in our trained models, the *aggregate* distribution of parameters was very similar between trained and untrained models, again highlighting the difficulty of constructing task-performing networks from bottom-up considerations alone.

Third, in addition to training such models, Jaxley will also enable numerous other applications: We showed how backprop enabled gradient-based Bayesian inference (Fig. 2h) [118, 119], and that it allowed us to investigate the stability of dynamical systems by computing their Lyapunov exponents (Fig. 4b), and made it possible to study adversarial attacks on biophysical network models (Fig. 5h). One will also be able to use gradients to compute maximally excitable stimuli [120, 121], or to design optimally discriminative experiments [122].

A central challenge in training biophysical models to perform computational tasks will be to bridge the timescales between biophysical mechanisms (milliseconds) and behavior (seconds). We showed that it is possible to fit data of up to one second length (Fig. 2d), but optimizing models on tasks that require backpropagating gradients along even longer simulations is challenging. In addition, opening up the possibility to learn the parameters of biophysical models with virtually no limit on the number of parameters will make mechanistic models prone to overfitting. Highly flexible models will also be more likely to identify *some* parameter sets that fit the data well, even if the model is wrong. To mitigate these issues, one should carefully design the loss function, the methods for model comparison, and the evaluation of the predictive quality on unseen data. One approach can be, for example, to train ensembles of models from different initial conditions, and to subsequently investigate which model-properties are robustly preserved across the ensemble [16].

Jaxley contributes to a growing body of work on simulators for neuroscience, while offering key advantages for biophysically-detailed simulation. Like previous simulation toolboxes such as Neuron [105], G_ENESIS_ [32], or N_EST_ [31], Jaxley implements an implicit Euler solver required to solve the stiff dynamics of morphologically detailed biophysical neuron models (note that such a solver is not implemented in standard toolboxes for differential equations [52, 123]). Unlike these widely used neuroscience toolboxes, Jaxley can automatically differentiate through this solver, thereby allowing to perform gradient descent without having to build differentiable emulators of biophysical models [72, 124]. In addition, its just-in-time compilation [125] and GPU parallelization capabilities [36–38, 54, 126] allow for fast simulation and training and enable scaling to large networks and datasets. By building upon the framework JAX [49], Jaxley will further benefit from advances in the deep learning community at scaling and training large simulations. For example, for some tasks and training paradigms, forward mode automatic differentiation [127, 128] or evolutionary algorithms [42] have been reported to perform similarly to (or sometimes even better than) backprop. Jaxley directly supports GPU-accelerated implementations of either of these algorithms, opening up possibilities to develop new methods for training of biophysical neural systems. Jaxley has a flexible and easy-to-use interface and offers a growing and easily-extensible set of channels and synapses.

New experimental tools allow to measure connectivity [129, 130], morphology [131, 132], genetic identity [109], and activity [133] of neural circuits at increasing levels of detail and scale. Jaxley is a powerful new tool which allows to integrate measurements of connectivity and morphology into biophysical simulations, while allowing the resulting networks to be fitted to data or computational tasks—just like deep neural networks. This will enable investigations of the biophysical basis of neural computation at unprecedented scales.

## Acknowledgments

We thank Nathanael Bosch, Philipp Hennig, Sarah Müller, Seth Axen, Thomas Euler, Matthias Bethge, Thomas Zenkel, Federico D’Agostino, Jan-Matthis Lueckmann, and all members of our research groups for discussions. This work was supported by the German Research Foundation (DFG) through Germany’s Excellence Strategy (EXC 2064 – Project number 390727645) and the CRC 1233 “Robust Vision”, the German Federal Ministry of Education and Research (Tübingen AI Center, FKZ: 01IS18039A), the ‘Certification and Foundations of Safe Machine Learning Systems in Healthcare’ project funded by the Carl Zeiss Foundation, and the European Union (ERC, “DeepCoMechTome”, ref. 101089288, “NextMechMod”, ref. 101039115). Views and opinions expressed are however those of the authors only and do not necessarily reflect those of the European Union or the European Research Council Executive Agency. Neither the European Union nor the granting authority can be held responsible for them. MD, KLK, JB, MP, MG, and JKL are members of the International Max Planck Research School for Intelligent Systems (IMPRS-IS).

## Author contributions

Conceptualization, Methodology: MD, PB, PJG, JHM. Software and Investigation: MD, KLK, MP, JB, MG, ZH, JKL, CS. Analysis: MD, KLK, MP, PB, PJG, JHM. Writing: MD, KLK, MP, PB, PJG, JHM. Writing (Review & Editing): JB, ZH, MG, JKL, CS. Funding acquisition: PB, JHM. Supervision: PB, PJG, JHM.

## Methods

### Code availability

Our toolbox, Jaxley, is openly available at https://github.com/jaxleyverse/jaxley. Tutorials and examples of how to use the toolbox are available at https://jaxley.readthedocs.io. A collection of channels and synapses for use with Jaxley is available at https://github.com/jaxleyverse/jaxley-mech. All code to generate results and figures is available at https://github.com/mackelab/jaxley_experiments.

### Solving differential equations of biophysically detailed single neuron models

Below, we describe the numerical solver for the differential equations that define dynamics in morphologically detailed neuron models. We used an implicit Euler solver for the voltage equations, and an exponential Euler solver for the gates. We simulated voltage at internal nodes and, like Neuron, also simulated terminal nodes at branch points. To solve the tridiagonal system of equations of every branch, Jaxley allows to choose between the Thomas algorithm [134] and the algorithm presented by Stone [135]. We used Stone’s algorithm for all simulations.

We performed a leapfrog update of voltage equations and gate equations. As is also done in Neuron, at every time step, the current through mechanisms (channels and synapses) was evaluated twice at voltage values that differ by 0.001 mV. This allowed to infer the voltage-dependent and voltage-independent contributions to the current dynamics (which is required by the implicit Euler solver of the voltage equations). In order to achieve a high degree of parallelization, we modelled every branch in the cell (or network) with four compartments (with the exception of the network shown in Fig. 1f, for which we used two compartments). We split branches that were longer than 300 µm into sub-branches until each sub-branch was shorter than 300 µm.

### Robust training of biophysical models

We use the following tools to improve training accuracy and robustness. First, we used parameter transformations that ensure that optimization of bounded parameters can be performed in unconstrained space and that biophysical parameters are on the same scale. In particular, we used an inverse sigmoid transformation 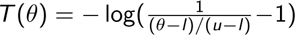, where *l* is the lower bound and *u* is the upper bound. Second, we used a variant of Polyak stochastic gradient descent [56, 136], which computes the step as step = 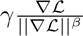. This optimizer overcomes large variations of the gradient between steps. We used values of *β* ∈ [0.8, 0.99]. In some applications, we further multiplied the gradient with the loss value L [56, 136], such that the optimizer automatically reduced the learning rate towards the end of training. Third, when necessary, we used different optimizers for different types of parameters. This affects the Polyak gradient descent optimizer because it normalizes by the gradient norm. Fourth, we performed multi-level checkpointing to reduce the memory requirements of backpropagation through time. Fifth, when we observed vanishing or exploding gradients, we performed truncated backpropagation through time. We used this only for the hybrid mechanistic/statistical model of the retina. When performed, we interrupted gradient computation every 50 ms. All of these tools are implemented in Jaxley.

### Training overview

In Table 1, we list the training procedure for all tasks, including the number of optimized parameters, the number of branches that the model has (note that each branch is modelled with 4 compartments), the compute device we trained on, the number of gradient steps we took to arrive at the model shown in the figures, and the compute time it took to perform one gradient step.

**Table 1.** Overview of tasks and their training procedure.

| Task | params | branches | device | gradient steps | time per step |
| --- | --- | --- | --- | --- | --- |
| L5PC synthetic (Fig. 2b) | 19 | 339 | CPU | 10 | 5 sec |
| Allen cell 1 (Fig. 2d, left) | 19 | 103 | CPU | 10 | 12 sec |
| Allen cell 2 (Fig. 2d, right) | 19 | 99 | CPU | 10 | 12 sec |
| Fit conductance profiles (Fig. 2f) | 1390 | 339 | CPU | 500 | 0.04 sec |
| Nonlinear neuron (Fig. 2g) | 72 | 3 | CPU | 6400 | 0.0017 sec |
| Hybrid retina model (Fig. 3) | 607 | 147 | GPU, A100 | 560 | 55 sec |
| RNN for evidence integration (Fig. 4) | 109 | 62 | CPU | 2000 | 12 sec |
| Delayed-match-to-sample task (Fig. 4) | 459 | 152 | GPU | 1000 | 36 sec |
| Biophysical network for MNIST (Fig. 5) | 105k | 2000 | GPU, V100 | 26k | 28 sec |

### Fitting layer 5 pyramidal cells to somatic voltage recordings

#### Active mechanisms

We used two types of sodium channels, five types of potassium, two types of calcium, and a hyperpolarization-activated channel and inserted them in soma, basal and apical dendrites, and axon as done by Van Geit et al. [39].

#### Parameter initialization, parameter sharing, and parameter constraints

To fit the synthetic data shown in Fig. 2b, we optimized 19 parameters. These were the same parameters as used by Van Geit et al. [39] but our model did not contain a persistent sodium channel in the axon (following Deistler et al. [137], which demonstrated that using a persistent sodium channel is inconsistent with experimental measurements).

We used the same parameter search bounds as Van Geit et al. [39] for the synthetic data, but we enforced that somatic potassium existed (lower bound 0.25 mS/cm^2^). For the experimental recordings from the Allen Cell Types Database, we used slightly larger bounds for two of the parameters to increase the flexibility of the model: We used an upper bound of 2 mS/cm^2^ (instead of 1 mS/cm^2^) and a lower bound of 10 ms for the delay of the calcium buffer (instead of 20 ms). We used a log-uniform distribution to initialize the delay of the calcium buffer. To fit recordings from the Allen Cell Types Database (IDs 485574832, 488683425, 480353286, and 473601979), we modified the leak conductance to 5·10^-5^, 10^-4^, 10^-4^, and 10^-4^ mS/cm^2^, respectively, leak reversal potential to −88, −88, −95, and −88 mV, capacitance to 2, 4, 3, and 2 µF/cm^2^, respectively, and potassium reversal potential to −70 mV.

#### Summary statistics

Optimizing parameters of biophysical models with gradient descent requires that the loss function, and therefore also the summary statistics, are differentiable. This is not the case for typically used features such as spike count, which led us to defining different summary statistics.

For the synthetic data (Fig. 2b), we split the voltage trace into two windows and used the mean and standard deviation of these two windows as summary statistics. This led to a total of four summary statistics. To standardize the data, we divided the mean voltages by 8.0 and standard deviations by 4.0.

To fit data from the cell-types database, we used a set of four such windows. We defined the first window as the 20 ms after stimulus onset and used the maximal voltage of this window as summary statistic. This avoids that the model spikes before the experimental data. We defined the second window as a 15 ms window around the first spike in the experimental voltage and used the maximum value within this window as summary statistic. The third and fourth window spanned the next 60 and 905 ms and we used the mean and standard deviation during these windows. To standardize the summary statistics, we divided maximal voltages by 50.0, mean voltages by 10.0, and standard deviations by 5.0.

#### Training procedure

We used our variant of Polyak gradient descent with *γ* = 1*/*3 and *β* = 0.8 for the synthetic data and the experimental data. For both tasks, we use a mean absolute error loss to standardized summary statistics. For models based on the Allen Cell Types Database, we performed five initial simulations and initialized gradient descent runs at the parameter set which had lowest loss (among those five runs).

For both experiments, we trained for ten optimization steps. We also ran the genetic algorithm (IBEA-DEAP as implemented in the BluePyOpt package, version 1.14.11 [39]) for ten iterations, with ten simulations per iteration. We ran all experiments on an Apple MacBook Pro M3 CPU. For the synthetic data, a single gradient step took around five seconds, for the experimental data (which had a longer simulation time, but the neurons had fewer branches), it took around twelve seconds.

To compute how many simulations were needed for the genetic algorithm and gradient descent to obtain good solutions (Fig. 2c), we thresholded the loss value (see figure captions) and computed the number of simulations (or total compute time) divided by the number of converged runs.

### Fitting voltage recordings of all branches

#### Active mechanisms

We used the same ion channels as for the task of fitting electrophysiology traces. To generate the observation, we sampled from a Gaussian process the conductance profiles as a function of the euclidean distance from the soma: we did this for the maximal conductances of all active ion channels in the apical dendrite (three active mechanisms) and the axon (seven active mechanisms).

#### Parameter initialization, parameter sharing, and parameter constraints

For the active mechanisms in the apical dendrite and the axon, we defined one parameter per branch in the cell and initialized parameters randomly and independently from each other. We used a single parameter for each mechanism in the soma and basal dendrite. Parameters had the same lower and upper bound as for the L5PC.

#### Training procedure

We used a mean absolute error loss function between the observed voltages and the simulated voltages, evaluated at every fifth time step between 1 ms and 5 ms of simulation time. We used our custom optimizer with *β* = 0.8, momentum 0.9, and learning rate 0.1. We trained the system for 500 iterations. We regularized the optimization such that neighboring branches had similar conductance values. In particular, we added to the loss the term 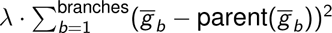, with regularization strength *λ* = 0.001.

One iteration of gradient descent took 0.04 seconds on an Apple MacBook Pro CPU.

#### Bayesian inference

To perform Bayesian inference over the parameters, we used a uniform prior over the free parameters with equivalent bounds as with the original training procedure. We also incorporated the same regularizer into the prior distribution. Since the optimization is performed in unconstrained space, we also conducted MCMC in this unconstrained space through a change of variables. We utilized a Laplace log-likelihood with a scale parameter of *λ* = 0.001, ensuring that the unnormalized posterior log density matched the loss function of the optimization problem.

We sampled from the posterior distribution with Hamiltonian Monte–Carlo (HMC) [138], which leverages the available gradient to efficiently sample from high-dimensional distributions. We used the BlackJAX [139] implementation of HMC. For each step, we performed five integration steps for the Hamiltonian dynamics with a step size of 0.01, leading to an average acceptance rate of ≈ 65% (which indicates good performance for HMC [140]).

We initialized 200 chains with samples from the prior distribution. Each chain was run for 500 iterations in parallel on a single NVIDIA GeForce A100 GPU, and only the last sample of each chain was considered.

To visualize the learned conductance profile, we discretized distance from soma into eleven bins and grouped all parameters within each bin. We then calculated histograms for each of the bins and generated a spline interpolation of all quantile lines (Fig. 2h).

### Nonlinear single neuron computation

#### Active mechanisms

The model contained sodium, potassium, and leak channels in all branches, with dynamics following the default implementation of the Neuron package [105].

#### Parameter initialization, parameter sharing, and parameter constraints

We trained sodium, potassium, and leak maximal conductances, as well as radius, length, and axial resistivity of every compartment in the model. Parameters were initialized randomly within uniform bounds. The bounds were [0.05, 1.1] for sodium, [0.01, 0.3] for potassium, and [0.0001, 0.001] for leak, all in mS/cm^2^. The bounds for the radius were [0.1, 5.0] µm, for the length [1, 20] µm per compartment, and for the axial resistivity [500, 5500] Ωcm.

#### Training procedure

We used a mean absolute error loss function between the simulated voltages after 3 ms of simulated time and the class label (35 mV or −70 mV). We used the Adam optimizer with learning rate 0.01, and batch size 1. We trained the system for 200 epochs. One iteration of gradient descent took 1.7 ms on an Apple MacBook Pro CPU.

### Hybrid model of the retina

#### Details on the data

Below, we describe the features of the experimental data which are most relevant to our training procedure (for full experimental details see Ran et al. [73]).

Images of 15 × 20 pixels were presented to a mouse retina. Each pixel had a size of 30 × 30 µm. Each image was presented for 200 ms, and a total of 1,500 images were presented. The images were centered on recording fields of the calcium activity. In total, 15 recording fields were made for the off alpha cell used in our work. Within each recording field, Ran et al. [73] defined a variable number of regions of interest, within which the calcium activity was recorded. In total, the data contained 232 regions of interest.

#### Data preprocessing

To denoise the calcium data, we lowpass-filtered the raw calcium data with a butterworth filter and a cutoff frequency of 7 Hz. We z-scored the resulting signal, with a different mean and standard deviation for each region of interest.

Next, we generated a single label for each image. We did this by performing a linear regression from image onto delayed calcium signals, and then used the calcium at the delay which was most predictive (i.e., had highest Pearson correlation between linear regression prediction and data). This led us to a delay of 1.8 seconds. As label, we used the low-pass filtered calcium value after this delay (starting from image onset).

#### Hybrid model

We modelled photoreceptors as a spatial linear Gaussian filter. The filter had a standard deviation of 50 µm. We modelled Bipolar cells as point neurons with a nonlinearity. The point neurons were spaced on a hexagonal grid with distance between neurons being 40 µm. The nonlinearity was taken from values measured by Schwartz et al. [74]. We connected every bipolar cell onto every branch of the retinal ganglion cell which was within 20 µm of the bipolar cell. Within every branch of the RGC, the synapse was made to the compartment which had the minimal Euclidean distance to the BC.

We computed the calcium activity of the model as the intracellular calcium value of the compartment that was the closest to the experimental recording site. We convolved this value with a double-exponential kernel to model dynamics of the calcium indicator. We used a rise-time of 5 ms and decay-time of 100 ms.

#### Active mechanisms in the retinal ganglion cell

Our model of the retinal ganglion cell contained six ion channels which were developed based on measurements from the retinal ganglion cells in cats [76]. The channels were a sodium channel, a leak channel, a delayed-rectifier potassium channel, a transient potassium channel, a calcium-dependent potassium channel, and a calcium channel. The model had all of these channels in every compartment of the model.

#### Parameter initialization, parameter sharing, and parameter constraints

We initialized all membrane conductances at previously published values [76], with an exception of sodium conductances which we initialized at 0.15 in the soma and 0.05 in the dendrite. We sampled the initial synaptic strengths randomly within 0 and 0.1 nS, and then divided the synaptic conductance by the number of postsynaptic connections that a bipolar cells makes (such that, in expectation, every BC has the same impact on the RGC). We initialized the axial resistivity of every compartment at 5,000 Ωcm. Finally, we initialized the radius of every dendritic compartment at 0.2 µm. We kept the diameter of the soma constant at 10 µm.

We trained the following set of parameters: One value for each maximal conductance in the soma (six parameters), one value for each maximal conductance in the dendrites, shared across all dendrites (six parameters), one value for each branch radius (147 parameters), one value for the axial resistivity of each branch (147 parameters), and one value for each synaptic conductance from the bipolar cells onto the retinal ganglion cells (250 parameters). In total, the model had 556 parameters.

We used the following bounds for optimization: For somatic conductances we used [0.05, 0.5] for sodium, [0.01, 0.1] for potassium, [10–5, 10–3] for leak, [0.01, 0.1] for transient potassium, [2·10^-5^, 2·10^-4^] for calcium dependent potassium, and [0.002, 0.003] for calcium. All membrane conductance units are mS/cm^2^. For dendritic conductances, we used the same bounds apart from the lower bound of 0 for sodium. For the branch radii, we used bounds of [0.1, 1.0] µm [73]. For the axial resistivities, we used [100, 10,000] Ωcm. For the synaptic conductances, we used [0.0, 0.2] nS.

#### Training procedure

We trained the model with Polyak stochastic gradient descent with momentum. We considered every kind of parameter (somatic conductance, dendritic conductance, radii, axial resistivities, and synaptic conductances) as separate parameters (which influences the computation of the gradient norm in Polyak stochastic gradient descent). We first trained the model for ten epochs with a learning rate of 0.01 and a momentum of 0.5. We used *β* = 0.99 to compute the norm in Polyak stochastic gradient descent.

We used a batchsize of 256 and used two level checkpointing to reduce the memory of backpropagation. To avoid vanishing or exploding gradients, we truncated the gradient in time. Specifically, we reset the gradient to 0 after 50, 100, and 150 ms of simulations.

We trained the model with mean absolute error loss between the experimentally measured (lowpass filtered) calcium value and the model predicted calcium value after 200 ms. Recording sites for which no recording was available in the data were masked out in the loss computation.

#### Receptive field computation

We followed Ran et al. [73] to compute receptive fields. We used automatic smoothness detection (ASD) [141] with 20 iterations of evidence optimization. We standardized all receptive fields to range from 0 to 1 and, for contours, thresholded the receptive fields at a value of 0.6.

#### Inductive bias

To evaluate the inductive bias of the hybrid model, we trained several such models, each on a reduced dataset. We reduced the dataset by using only a subset of stimulus/recording pairs from one scan field. We repeated this procedure over seven scan fields and five sizes of stimulus/recording pairs, namely 32, 64, 128, 256, 512. We trained each model with stochastic gradient descent as described above. To ensure that the gradient is stochastic (which can improve learning [142]), we used batchsizes of 4, 4, 8, 16, and 32 for the five dataset sizes, respectively. We performed early stopping based on a validation set that contained 25 % of the training dataset and evaluated performance on 512 test datapoints. To avoid exceedingly long training times, we trained for at most 100 steps.

For the artificial neural network, we used a multi-layer perceptron with three hidden layers of [100, 100, 50] units and with ReLU activation functions.

### A recurrent neural network of biophysical neurons performing an evidence integration task

#### Active mechanisms

All recurrent units used the same ion channels as for the task involving the layer 5 pyramidal cell, analogously inserted in soma, basal and apical dendrite. The two readout neurons were passive. The synapses were conductance-based, as described by Abbott and Marder [88].

#### Parameter initialization and parameter constraints

The recurrent units were connected with a probability of 0.2 by synapses from the soma of the presynaptic neurons to the apical dendrite of the postsynaptic neurons. All recurrent units had synapses onto the two readout neurons. 50 % of recurrent units had inhibitory outgoing synapses, created by setting their synaptic reversal potential to −75 mV, and the other half were excitatory with a synaptic reversal potential of 0 mV. Initial values for the maximum synaptic conductances were drawn from a standard normal distribution scaled by an initial gain *g* such that the bulk of the eigenspectrum of the synaptic weights lay in a circle on the complex plane with radius *g* (after multiplying inhibitory synapse weights by −1) [143]. We presented inputs by stimulating neurons at their basal dendrite. We set *g* = 5.0*π /* 5 · 10^3^, which is close to the transition point between stable and chaotic dynamics. For recurrent connections, we set the rate constant for transmitter-receptor dissociation rate (the *k*_ parameter [65]), that influences the synaptic time-constants, to 1/1.0 ms. For connections to the rate-based readout neurons, we used slower synapses, with *k*_ set to 1/40 ms. All recurrent units received stimulus input scaled by random initial weights drawn from N (0, 0.1).

We trained the maximal synaptic conductances constrained to the range [0, 3 · max(*g*)]. We also trained the weights of the stimulus input to each neuron in the network restricted to the range [-0.2, 0.2]. The model had 109 trainable parameters.

#### Stimulus generation

Stimuli were generated by sampling values from normal distributions and then low-pass-filtering them with a maximum frequency cutoff of 2500 Hz. For training, we sampled values from N (*µ*_−_, 0.05) and N (*µ*_+_, 0.05), where *µ*_−_ ∼ N (−0.005, 0.0002) and *µ*_+_ ∼ N (0.005, 0.0002).

#### Training procedure

We used a batch size of four, three levels of checkpointing, the Adam optimizer with a learning rate of 0.01, 2000 gradient steps, and gradient normalization with *β* = 0.8. We sampled new training data at every gradient step. Sweeps were used to inform the choice of hyperparameters. We used a cross entropy loss function with logits calculated as the mean readout activities in the last 20 ms of stimulus presentation. The training time was 7 hours on an Intel(R) Xeon(R) Silver 4116 CPU @ 2.10 GHz.

### A recurrent neural network of biophysical neurons performing a delayed-match-to-sample task

#### Active mechanisms

We used the same channel mechanisms, compartments and synapses as in the evidence integration task.

#### Parameter initialization and parameter constraints

We used a network of 50 units (in line with previous work [47]), with a recurrent connectivity probability of 0.05, and all units connecting to the readout neurons. 80 % and 20 % of the neurons had excitatory and inhibitory outgoing connections, respectively. We set *g* = 5.0*π /* 5 · 10^3^. We set the *k*_ parameter of recurrently connected neurons to 1, and to 0.1 for the slower synapses onto the readout neurons. The connection probability from input to (the basal dendrite of) recurrent units was 0.1, with initial weights drawn from U[0,1].

We trained the maximal synaptic conductances, constrained to the range [0, ∞), using a SoftPlus function. We also trained the weights of the stimulus input to each neuron in the network, restricted to the range [0, 4], as well as *k*_ restricted to [0.05, 2]. This resulted in 459 parameters.

#### Stimulus generation

Stimuli consisted of square pulses with additive Gaussian noise sampled from N (0, 0.001). The onset period was 20 ms, and the stimulus and response durations were 50 ms. The delay period changed throughout the training procedure. Initial delay durations were drawn uniformly from U[50,150]. The average delay duration was increased in steps of 100 ms, whenever the network got at least 95 % accuracy on a single batch, till U[450,550] was reached.

#### Training procedure

We used a batch size of 64, two levels of checkpointing, the Adam optimizer with a learning rate of 0.001, which was exponentially decayed to 0.0001 over 1000 epochs. We used a per time step cross entropy loss function, computed during the response period of the task. The training time was around 10 hours on an NVIDIA GeForce RTX 3090.

### Computing Lyapunov exponents

We can quantify the stability of recurrent networks by measuring the average rate of divergence or convergence of nearby trajectories, which is given by the maximal Lyapunov exponent. To obtain the maximal Lyapunov exponent, we first discretised our model to obtain **x***_t_*_+1_ = **f**(**x***_t_*), where **x***_t_*is a vector of all dynamic variables (e.g., voltages, gating variables) at time *t*, and **f** is one step of the chosen solver. We can then define the maximal Lyaponuv exponent as: 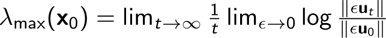 where **u**_0_ is a perturbation to the initial state of the system **x**_0_. We measured the evolution of infinitesimal perturbations to **x**_1:*T*_ by calculating the Jacobian at each point along a trajectory.

We used the following numerical algorithm to approximate *λ*_max_(**x**_0_) [90, 91]: First, we generated an initial state **x**_0_ and initial unit norm vector **q**_0_. After discarding initial transients for 4 seconds of simulation, we let the system **x***_t_*_+1_ = **f**(**x***_t_*) and **q***_t_*_+1_ = *D***f**|**_x_ q***_t_* (where *D* denotes the Jacobian) evolve for *T* = 2.4 · 10^5^ timestep (a further 6 seconds). Note that **q***_t_*_+1_ can be efficiently computed using Jacobian vector products in JAX [49]. At every timestep we computed *r_t_* = ∥**q***_t_*∥ and renormalised: 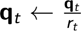. The maximal Lyaponuv exponent was then given by 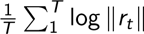.

The RNNs in Fig. 4b had 50 units with a 80% excitatory / 20% inhibitory split, and a connection probability of 0.2.

### A biophysical network that performs computer vision tasks

#### Active mechanisms and synapses

The first layer consisted of 28 × 28 neurons, each stimulated by a step current whose amplitude was proportional to a pixel value. Each neuron in the first layer had a ball-and-stick morphology: Each cell consisted of four compartments, where one compartment (the soma) had a radius of 10 µm and all other compartments had a radius of 1 µm and a length of 10 µm. The input and hidden layer contained sodium, potassium, and leak channels in all branches, with dynamics following the default implementation of the Neuron package [105]. The output layer consisted of ten neurons with ball-and-stick morphologies (like the input layer neurons) and with leak dynamics. We used conductance-based synapses as described by Abbott and Marder [88]. The layers of the network were densely connected. We set the synaptic rate constant for transmitter-receptor dissociation to *k*_−_ = 1*/*4.

#### Parameter initialization, parameter sharing, and parameter constraints

We optimized sodium, potassium, and leak maximal conductances of every branch in the network (50k parameters) and all synaptic conductances (50k parameters). We used the following bounds for the parameters: [0.05, 0.5] for sodium, [0.01, 0.1] for potassium, [0.0001, 0.001] for leak, [-5 / 28^2^ / 25, 5 / 28^2^ / 25] for synapses from the input to the hidden layer and [-5 / 64 / 25 / 2, 5 / 64 / 25 / 2] for synapses from the hidden layer to the output layer. We initialized sodium maximal conductances at 0.12 mS/cm^2^, potassium at 0.036 mS/cm^2^, and leak at 0.0003 mS/cm^2^. We initialized synaptic conductances as samples from a Gaussian distribution with mean 0 and standard deviation 1 / 28^2^ / 25 for the first layer and standard deviation 1 / 64 / 25 / 2 for the second layer.

#### Training procedure

We used a batch size of 16 and cross entropy loss based on the values (*v* + 65)*/*3, where *v* is the somatic voltage of the output neurons after 10 ms of simulation. We used a cosine learning rate schedule and trained the network for seven epochs. Each gradient step took 25 seconds on a V100 GPU.

#### Adversarial attacks

To perform the adversarial attacks, we performed optimization of the input with gradient descent. We normalized every gradient step and used a learning rate of 5.0. We used bounds of [0, 1] for all pixel values during optimization. We used a cross entropy loss function. We chose the target label for the adversarial attack randomly, but ensured that the target label is not the true label. We computed accuracy based on 128 adversarial attacks.

## Supplementary figures

**[Figure S1].**
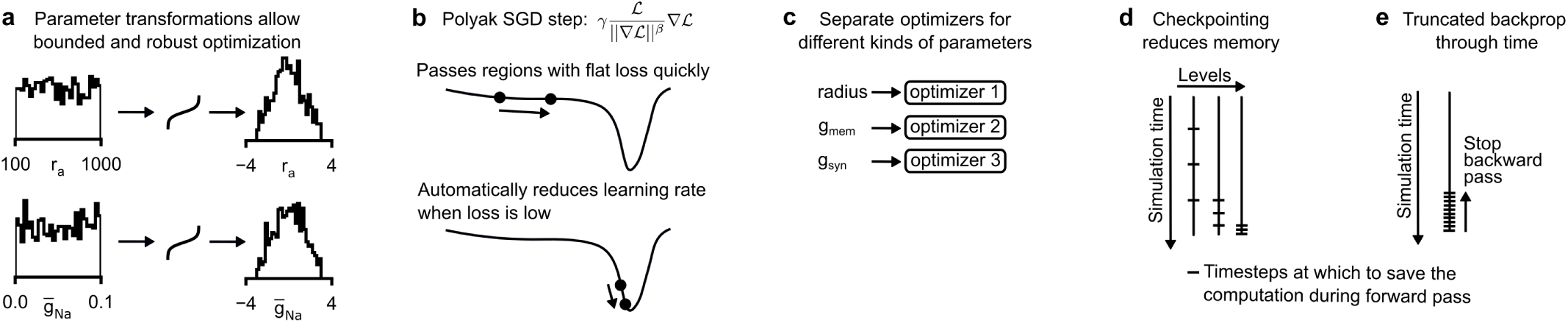
Robust and efficient gradient descent. (a) Histogram of two biophysical parameters (rows, random samples within optimization bounds) before (left) and after (right) parameter transformations. The transformation is designed such that the parameters are unconstrained and of the same scale. (b) Illustrative loss surface and update step made by our variant of Polyak gradient descent [56]. (c) We use different optimizers for different kinds of parameters. (d) Illustration of multi-level checkpointing [55]. We use checkpointing to overcome memory limitations of backpropagation of error. Multi-level checkpointing requires multiple forward passes per gradient computation, but typically reduces memory requirements. (e) Illustration of truncated backpropagation through time [144]. Truncated backpropagation through time allows to overcome vanishing or exploding gradients at the cost of providing only an approximate gradient.

**[Figure S2].**
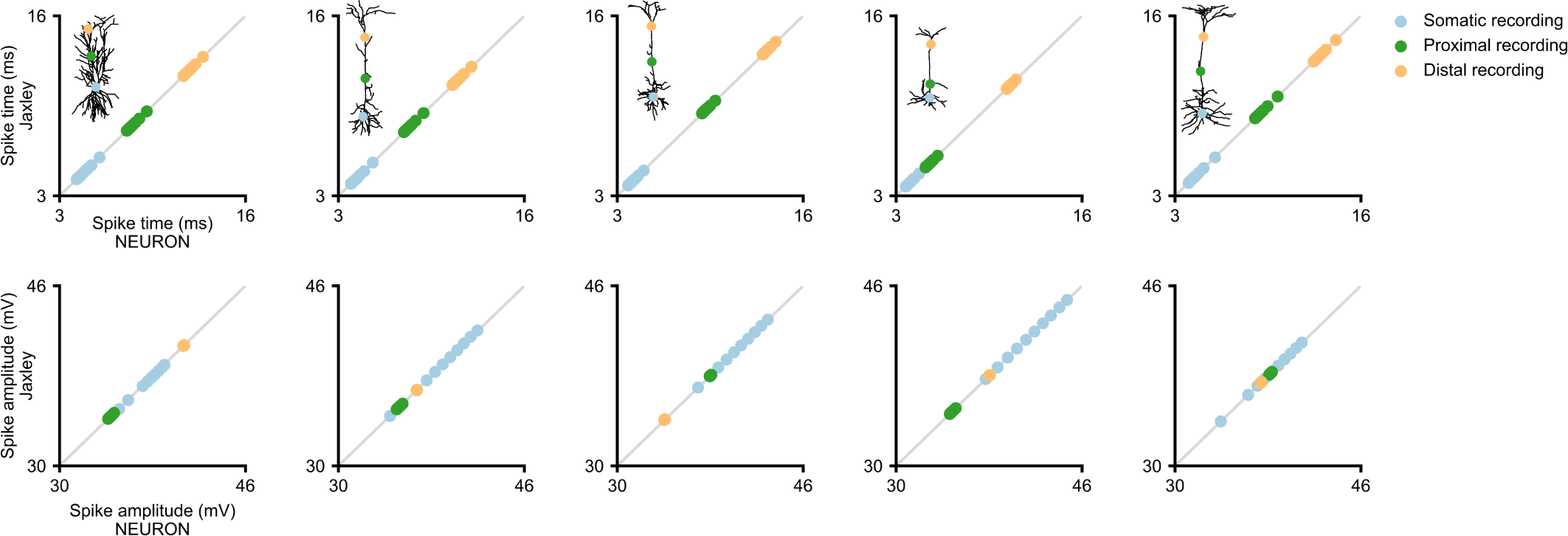
Accuracy of the implicit solver in Jaxley. Top: Spike time at three recording sites (somatic, proximal, distal) for input stimuli of 10 different amplitudes, ranging from 0.2 nA to 1.1 nA (individual dots). Columns are different morphologies from the Allen Cell Types Database. Bottom: Same as top, for spike amplitude.

**[Figure S3].**
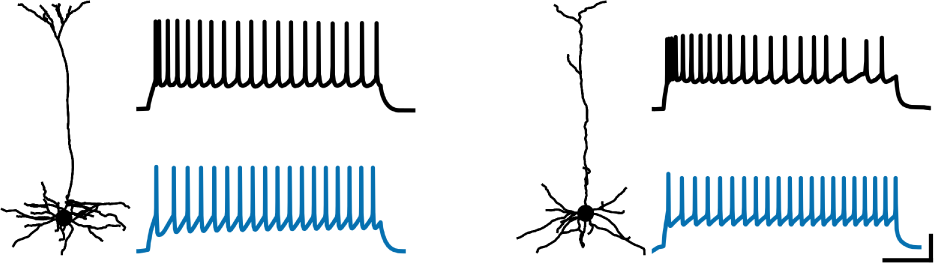
Gradient descent fits to recordings from the Allen Cell Types Database. Scalebars: 200 ms and 30 mV.

**[Figure S4].**
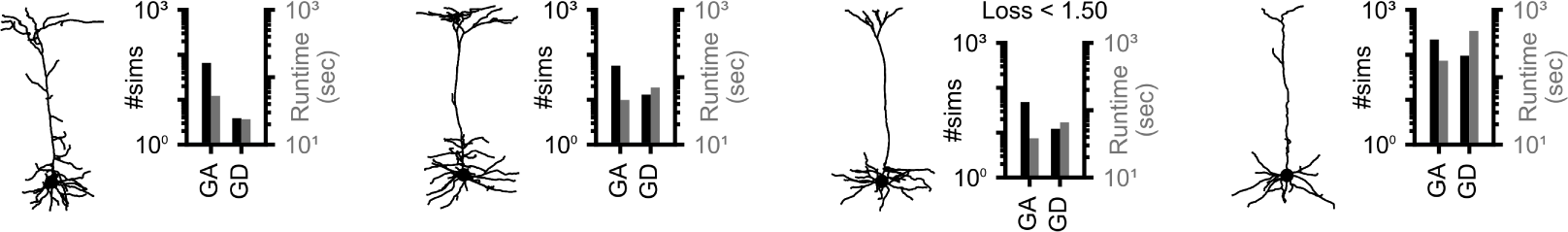
Fitting models to recordings from the Allen Cell Types Database. Number of simulations and runtime to reach a loss value of 1.5 or lower.

**[Figure S5].**
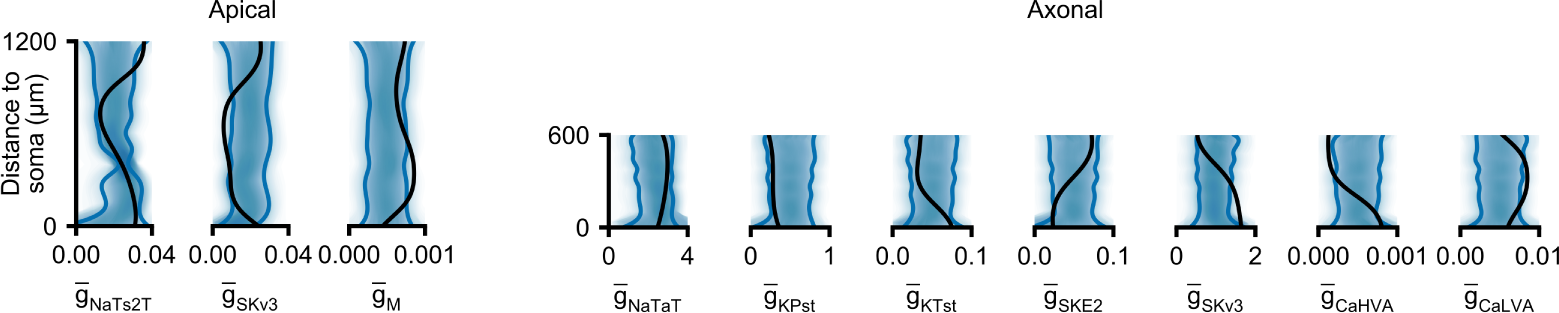
Bayesian inference of membrane conductances. We used Jaxley with gradient-based Hamiltonian Monte–Carlo to infer the posterior distribution over membrane conductances of a layer 5 pyramidal cell. Blue lines are 90 % confidence intervals, black line is the ground truth that was used to generate the synthetic observation.

**[Figure S6].**
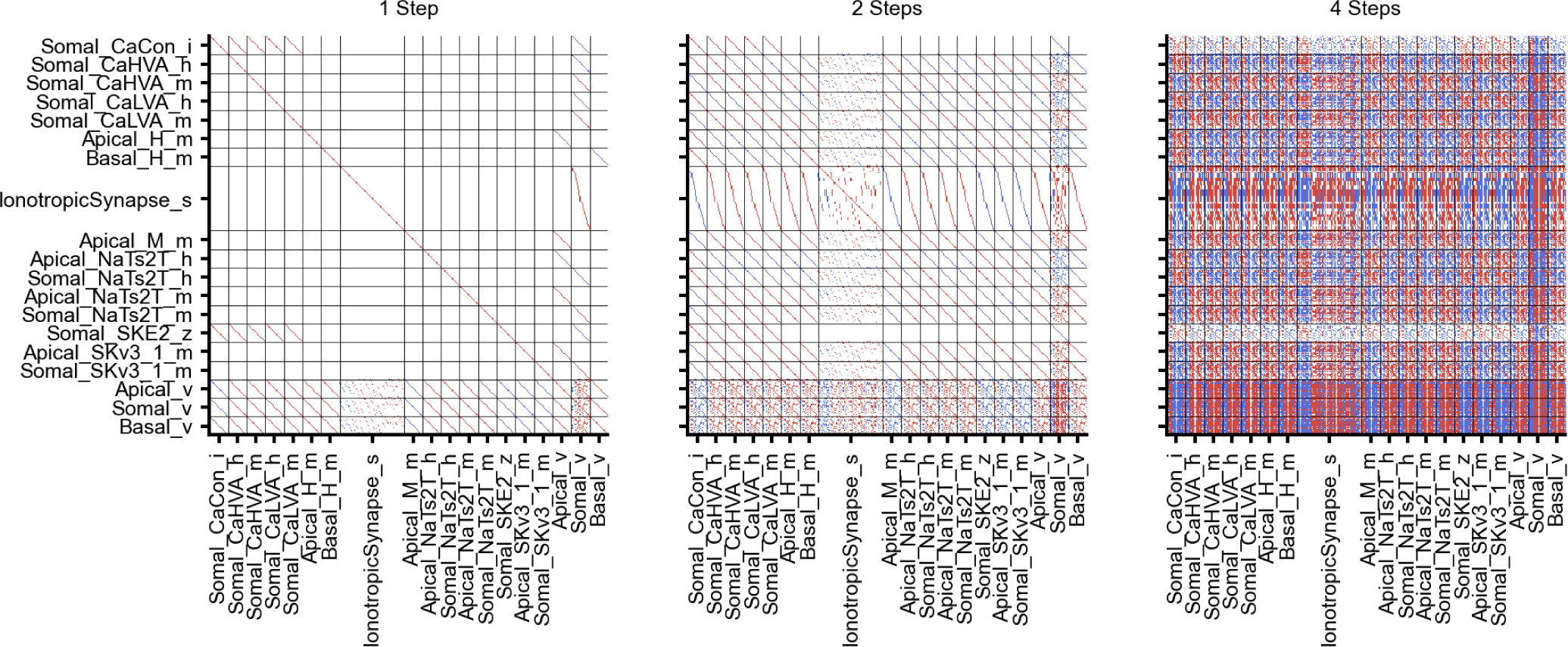
Jacobians reveal interactions between states Jaxley allows us to compute Jacobians of biophysical networks by using automatic differentiation. Here we show *D*f|_x0_, where *D*f denotes the Jacobian with respect to f, and f is 1, 2, and 4 steps of simulation with initial state x_0_. We used a recurrent network similar to those used in Fig. 5 (here: 20 units of which 5 are inhibitory; connection probability 0.2). As the scale between different states can be very different, we here just show the sign (red is positive, white is zero, and blue is negative).

**[Figure S7].**
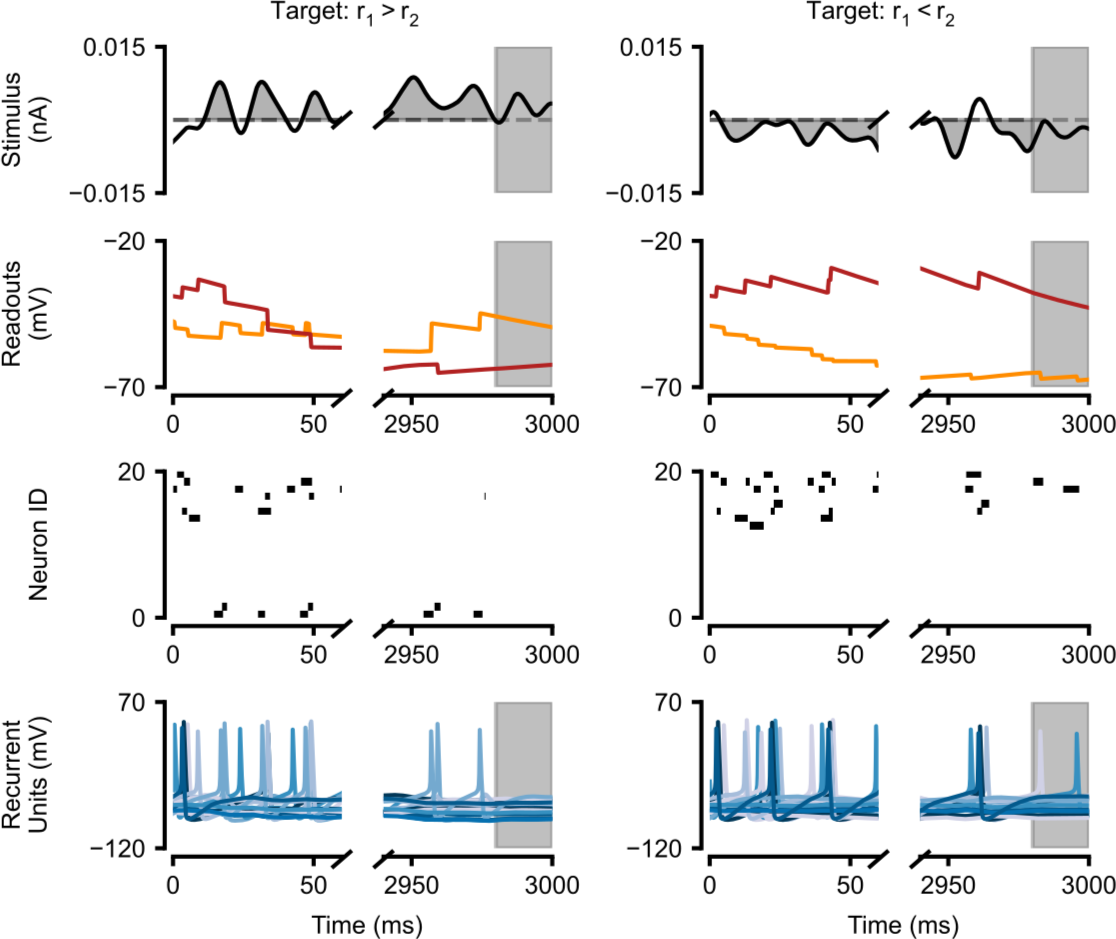
Generalization of the evidence integration task. Evidence integration task performance on three seconds of stimulus with positive integral (left) and negative integral (right). We used the same network parameters as in Fig. 5.

**[Figure S8].**
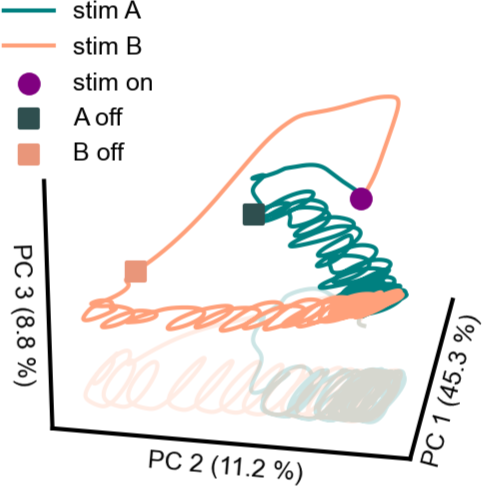
Long-term dynamics reveal transient coding. We here studied the long-term dynamics of the network from Fig. 5 trained to perform the delayed-match-to-sample task, in order to gain mechanistic insight into the model [101]. The figure shows the dynamics of the full state of the model (including ion concentrations), projected into principal component space, after briefly presenting either stimulus. We first smoothed each dynamic variable with a Hann window of 40 ms, and normalised, before computing the principal components. We find that, after stimulus offset, the trajectories first stayed separated, allowing the stimulus identity to be maintained for a duration that was long enough to bridge the delay period of 500 ms. Eventually, both trajectories ended on the same attractor and information about stimulus identity is lost.

**[Figure S9].**
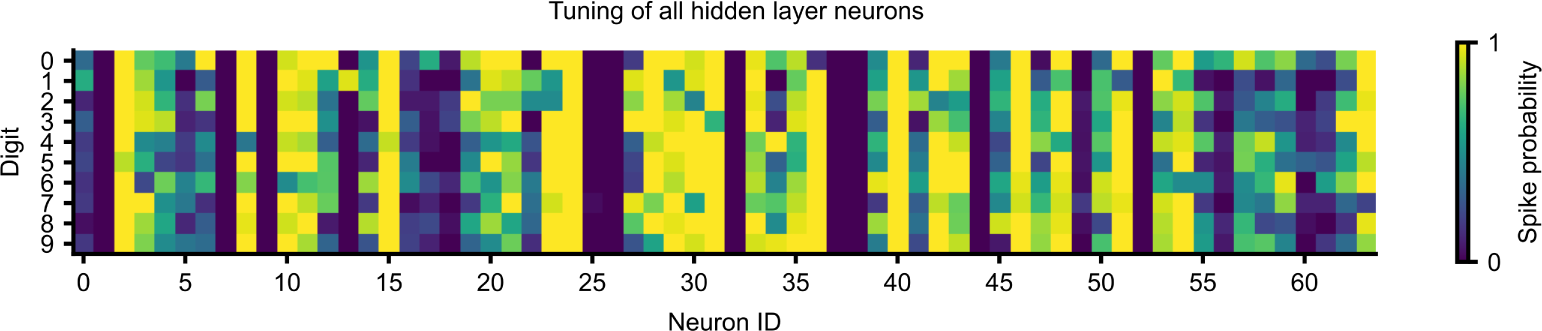
Hidden layer tuning of all neurons after training. We evaluated the fraction of images in the dataset for which each hidden neuron spiked.

**[Figure S10].**
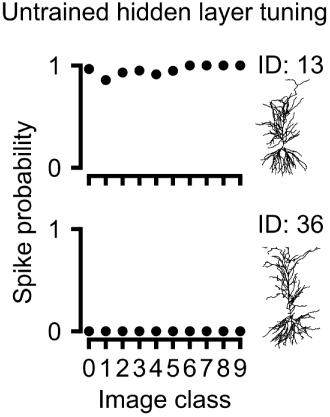
Hidden layer tuning before training. Before training, the two shown cells were untuned, but they developed tuning to specific digits (ON-tuning for digit ‘1’ for ID 13 and OFF-tuning for digit ‘0’ for ID 36) after training (Fig. 5g).

## Footnotes

1 Jaxley is openly available at https://github.com/jaxleyverse/jaxley

